# The role of white matter myelin in structural-functional network coupling

**DOI:** 10.1101/2025.07.24.666667

**Authors:** Mark C. Nelson, Wen Da Lu, Ilana R. Leppert, Heather A. Hansen, Christopher D. Rowley, Bratislav Misic, Christine L. Tardif

## Abstract

The brain is a complex network of neuronal populations interconnected by white matter tracts. The composition of these white matter connections (SC) shapes inter-regional signaling dynamics giving rise to spatial patterns of synchronous functional connectivity (FC). Several modeling approaches have proven useful for studying the mechanisms underlying the relationship between SC and FC. However, the myelination of white matter tracts — a major component of white matter connectivity — is not accounted for by conventional SC networks, and thus, has largely been excluded from models of FC. Here, we expand structure-function brain modeling by integrating a multi-feature white matter SC network. We use multi-modal MRI to compute an SC network with connections (edges) weighted by the caliber, myelination, and length of white matter tracts. We investigate the relationship of this multi-feature SC network with both haemodynamic and electromagnetic FC. Edge myelin was strongly predictive of FC in a pattern that was heterogeneous across brain regions and timescales of neural function. Edge myelin showed strong, frequency-specific interactions with both edge caliber and length suggesting a modulatory role for white matter myelin in structure-function coupling. This was further supported by antagonistic gradients of white matter myelin and structure-function coupling along the sensory-association axis. We describe in detail the individual and joint relationships between these major white matter features and multi-frequency FC. These results illustrate the advantage of a more comprehensive characterization of white matter in structure-function models and establish how white matter myelin — known for roles in conduction velocity, plasticity, and metabolic support at the microscale — shapes brain function at the macroscale.

## Introduction

Be it an ecosystem, an economy, or an encephalon, the form and function of complex systems are intricately linked(M. E. J. Newman, 2003). By applying the principles of network science(M. Newman, 2010) to the human brain(Bassett & Sporns, 2017; Fornito et al., 2016; Sporns, 2010; Suárez et al., 2020), a modular and small-world architecture is revealed which facilitates both specialized processing within regions, as well as efficient global communication. A fundamental component of this relationship is the arrangement and properties of the physical connections (edges) between regions (nodes), which constrain and shape the landscape of dynamic functional interactions(Sporns, 2011). The characterization of the mechanisms through which brain structure and function interact is a central goal in neuroscience.

Through the non-invasive lens of multi-modal magnetic resonance imaging (MRI), networks of macroscale connectivity between nodes can be estimated in the human brain *in vivo*. Functional connectivity (FC)(Salvador et al., 2005) typically quantifies the cross-correlation of blood-oxygen-level dependent (BOLD) functional MRI (fMRI) time series(Biswal et al., 1995), and structural connectivity (SC)(Hagmann et al., 2008; Honey et al., 2009) reconstructs the interconnecting white matter pathways by combining diffusion-weighted MRI with streamline tractography(Basser et al., 1994; Mori & Barker, 1999; Mori & Van Zijl, 2002). At this scale, network nodes amount to volumes of roughly 10^2^-10^4^ mm^3^ containing as many as 10^8^ neurons(Azevedo et al., 2009; Goriely, 2025), and structural edges likely represent the aggregate of both mono- and poly-synaptic white matter fiber pathways between these large gray matter regions.

While this spatial resolution may seem quite coarse, multiple modeling approaches have consistently demonstrated that SC based on diffusion MRI is predictive of BOLD-FC(Seguin et al., 2023; Suárez et al., 2020), and the link between them has implications for human behavior(Preti & Van De Ville, 2019; Seguin et al., 2020) and cognition(Litwińczuk et al., 2022; Popp et al., 2024; Shine et al., 2019). A spatially heterogeneous pattern of structural-functional network coupling has emerged(Liu et al., 2023; Preti & Van De Ville, 2019; Vázquez-Rodríguez et al., 2019; Zamani Esfahlani et al., 2022), which is closely aligned with the primary axis of cortical organization: the sensory-association (S-A) axis(Huntenburg et al., 2018; Sydnor et al., 2021). This places structure-function coupling alongside other large-scale features such as cortical hierarchy(Margulies et al., 2016; Mesulam, 1998), as well as finer-grained cortical features, including laminar differentiation(Paquola et al., 2019), intracortical myelin(Glasser & Van Essen, 2011; Huntenburg et al., 2017), neurotransmitter receptor distributions(Hansen et al., 2022) and gene expression(Burt et al., 2018).

Macroscale FC networks can also be derived from time series recorded with magnetoencephalography (MEG), which facilitates the assessment of neural function at higher frequencies (1-100+Hz) than BOLD-fMRI (0.01-0.5Hz). The fMRI signal is thought to reflect a superposition of these neurophysiological rhythms(Shafiei et al., 2022) and their cross-frequency interactions(Tewarie et al., 2016). Cross-frequency coupling plays an important role in the formation of brain networks both during rest(Florin & Baillet, 2015; Tewarie et al., 2016) and task(Brookes et al., 2016). Higher frequencies tend to reflect bottom-up sensory processing and lower frequencies top-down modulation(Baillet, 2017; Fries, 2015; Hillebrand et al., 2016; Scheeringa & Fries, 2019; Von Stein et al., 2000), which manifests as a gradient of timescales of neural function(Chaudhuri et al., 2015; Mahjoory et al., 2020; Murray et al., 2014) aligned with the S-A axis. Although these relationships –– as with BOLD-fMRI –– have been linked to multi-scale cortical structure(Helbling et al., 2015; Hunt et al., 2016; Mahjoory et al., 2020; Scheeringa & Fries, 2019) and conventional SC(Garcés et al., 2016), much less is known about the coupling between white matter structural and MEG-functional networks.

Thus, we come to a principal limitation of structure-function modeling: current models lack the requisite biological detail to draw strong conclusions about the underlying mechanisms(Suárez et al., 2020). One way to address this at the whole-brain level *in vivo* is by weighting the edges of structural networks using MRI-derived measures of white matter structure relevant to brain function(Assaf et al., 2008; Boshkovski et al., 2021; Caeyenberghs et al., 2016; Deligianni et al., 2016; Gast & Assaf, 2024; Lu et al., 2025; Mancini et al., 2018; Messaritaki et al., 2021; Nelson et al., 2023; Stikov et al., 2015; M. P. van den Heuvel et al., 2010). In particular, MRI-based estimates of white matter myelin density have been linked to both brain organization and function(Sonya Bells et al., 2017; Caeyenberghs et al., 2016; Gong et al., 2024; Messaritaki et al., 2021; Nelson et al., 2023; Petkoski et al., 2023).

The myelination of axonal fibers helps to regulate the timing of neural signaling(Huxley & Stämpfli, 1949; Lillie, 1925; Tasaki, 1939) supporting synchronization across distributed neural circuits(Nunez et al., 2015; Nunez & Srinivasan, 2014; Pajevic et al., 2014). Conversely, the plastic structure of myelin is itself shaped by neuronal activity, thus establishing a reciprocal mechanistic link between the modulation of white matter myelin and inter-regional functional coherence(Bacmeister et al., 2022; Chorghay et al., 2018; Douglas Fields, 2015; Fields et al., 2015; Mount & Monje, 2017; Zatorre et al., 2012). Although its role in conduction velocity is less clear at the macroscale(Berman et al., 2019; Mancini et al., 2021; Petkoski et al., 2023; Talidou & Lefebvre, 2025), white matter myelin is believed to support brain function through varied contributions including: gain control, circuit integration, resilience, and homeostasis(Karimian et al., 2019; Noori et al., 2020; Park & Lefebvre, 2020), as well as metabolism(Asadollahi et al., 2024; Nave, 2010; Ramos-Cabrer et al., 2025), and energetics(Harris & Attwell, 2012).

Myelination patterns in both the white matter and cortex are linked to the major S-A axis of brain organization. Along this gradient, intracortical myelin content decreases(Glasser et al., 2014; Glasser & Van Essen, 2011), and white matter tracts are myelinated progressively later(Fuster, 1995, 2008): primary sensory regions with high intracortical myelin myelinate their axons first. There is also evidence of both axis-general and axis-specific patterns of development for the strength of white-matter connectivity. Within-network connections at both ends of the axis become stronger over development, increasing network modularity and sharpening the differentiation of cortico-cortical systems. At the same time, inter-modular integration increases—but this occurs selectively through the strengthening of long-range association-hub connections, rather than uniformly across the axis(Sydnor et al., 2021). This topological pattern – – also supported by MRI-based metrics of white matter myelin(Boshkovski et al., 2021) –– is fundamental to the brain structure-function relationship(Sporns, 2010, 2011, 2013). Thus, a coordinated blueprint of cerebral myelination gives rise to a global axis of myelin content, which is linked to the organization of both brain structure and function.

Despite these links to brain function, white matter myelin has largely been excluded from structure-function models. How does white matter myelin relate to global FC, and does this relationship vary across brain regions and functional timescales? Here, we address these questions by integrating white matter tract features including myelin into a network model of FC. Specifically, we combine a network analysis approach with multi-modal MRI to quantify three major features of white matter connectivity at the network edge level: caliber, myelin, and length. We then use these features in an edge-wise multi-linear model to predict haemodynamic and electromagnetic FC. Based on previous work(Nelson et al., 2023), we expect that myelin and BOLD-FC will be inversely related globally. We further hypothesize that the regional coupling of myelin and BOLD-FC will be stronger, heterogeneous, and aligned with the S-A axis, in line with results using spectral(Preti & Van De Ville, 2019) and communication features(Liu et al., 2023; Vázquez-Rodríguez et al., 2019; Zamani Esfahlani et al., 2022) from conventional SC. Anticipating the coupling between myelin and MEG-FC is challenging, as few or no studies have explored this. However, it is reasonable to expect stronger, heterogeneous, and frequency-specific coupling regionally, as shown for conventional SC(Liu et al., 2023).

## Results

Multi-modal MRI data acquired at 3 tesla in 30 healthy adults (14 men, 16 women; 29±6 years of age) were used to estimate the SC and in-sample FC networks. Structural connections were derived by applying streamline tractography to diffusion MRI data, then weighted by edge-level estimates of the caliber, myelin, and length of white matter tracts. Edge caliber quantifies the total cross-sectional area of the intra-axonal compartment. Edge myelin quantifies myelin density and was derived from tractometry of magnetization transfer saturation images. The in-sample FC network was estimated as the zero-lag Pearson cross-correlation of node-wise time series from resting-state (task-free) BOLD-fMRI data (**BOLD_in_**). Group-averaging resulted in four connectivity networks (caliber, myelin, length, and BOLD_in_) from in-sample data. An additional seven out-of-sample FC networks were obtained in fully preprocessed(Shafiei et al., 2022) and group-averaged format: one from resting-state fMRI (**BOLD_out_**) and six from resting-state MEG corresponding to the canonical electrophysiological frequency bands: **delta** (2-4Hz), **theta** (5-7Hz), **alpha** (8-12Hz), **beta** (15-29Hz), **gamma_lo_** (30-59Hz), and **gamma_hi_** (60-90Hz). These were derived from data in healthy participants (n = 33; age range 22 to 35 years) made openly available through the Human Connectome Project (HCP; S900 release(Van Essen et al., 2013)). Thus, eight unique FC networks were separately modeled as linear combinations of three weighted SC networks including interaction terms for both caliber and length with myelin (**Figure 1a**). Modeling was performed at three spatial scales: global (whole-brain), pairwise combinations of resting-state networks, and at the level of individual nodes. Network nodes were defined using the Schaefer-400(Schaefer et al., 2018) cortical atlas. The correlation between all structural features and all FC networks is shown in **Figure S9**

**Figure 1.**
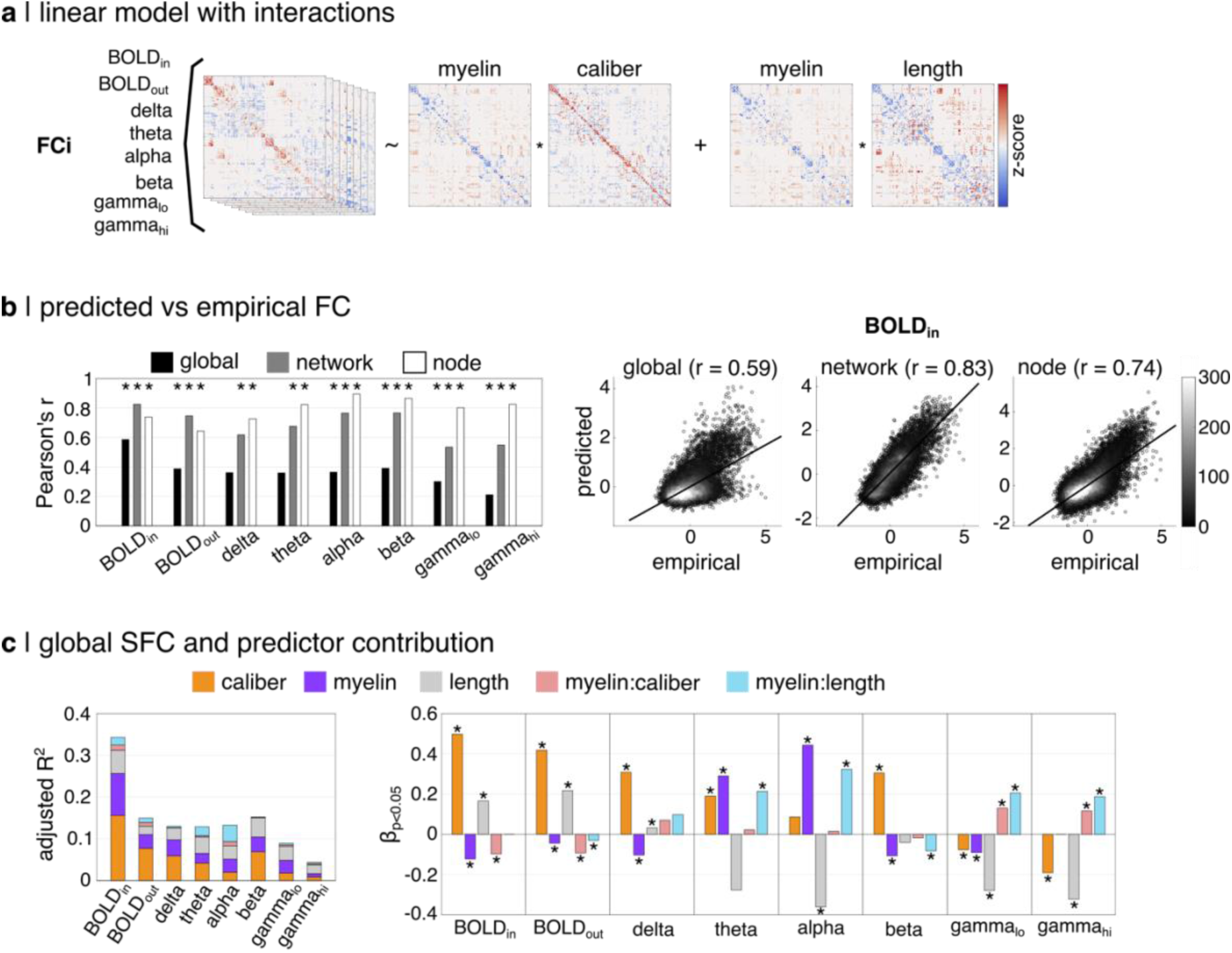
Relating white matter structural features to haemodynamic and electromagnetic functional connectivity. **(a)** An edgewise multi-linear regression model predicting FC from SC. FC was quantified using BOLD-fMRI and multi-band MEG. SC was quantified as the caliber, myelin and length of white matter tracts from multi-modal MRI. Interaction terms with myelin were included for caliber and length. All data was z-scored before modeling. **(b)** Pearson’s correlations of predicted vs empirical FC at global, network and node-level resolutions. Edges were pooled for correlations at network and node-levels. All correlations were significant (p_perm_ < 0.05), except global delta and theta. Scatter plots (right) are shown for BOLD_in_. **(c)** Global SFC strength (left) is quantified using adjusted R^2^. Colored bar segments show ratio of predictor contributions from dominance analysis. Standardized ß-coefficients (right) surviving parametric t-tests (p < 0.05) are shown. Asterisks additionally indicate significance from permutation testing (p_perm_ < 0.05) and 0 ∉ 95% confidence intervals.

### White matter myelin supports haemodynamic and electromagnetic connectivity

FC values predicted by the model correlated strongly with empirical FC data (**Figure 1b**). The correlation magnitude increased with model resolution for both MEG and fMRI, however the optimal resolution was modality-dependent and mirrored their temporal resolution differences: strongest at the network level for BOLD (r > 0.75, p_perm_ = 0.001) and at the node level for MEG (r > 0.70, p_perm_ = 0.001). Node-level alpha and beta models showed the highest correlations (r > 0.85, p_perm_ = 0.001). Delta and theta models failed to reach significance at the global level when applying a spatially-constrained permutation test.

We assessed the global relationship between white matter edge features and FC by pooling all upper triangle edges with SC ≠ 0 into a single model (**Figure 1c**). The strength of structure-function coupling (SFC) was inferred from model fit (adjusted coefficient of determination; R^2^). The contribution of each structural feature to the prediction of FC was quantified in two ways: (1) their relative importance from *dominance* analysis(Azen & Budescu, 2003; Budescu, 1993), and (2) the nature of their *association* from the sign and magnitude of standardized ß coefficients.

SFC was strongest for BOLD_in_ (R^2^ = 0.34), lowest for the gamma bands (R^2^ < 0.09), and roughly equivalent for BOLD_out_ and delta-beta (R^2^ ∼ 0.13-0.15). Myelin was a strong predictor of FC in both BOLD and MEG data. Myelin ranked 2^nd^ in predictor dominance behind caliber for BOLD_in/out_ and delta, dropped to 3^rd^ for theta and beta as the relative importance of length increased in MEG, and shared dominance with length for alpha and gamma as caliber decreased in importance at higher frequencies. The ß-coefficients for all predictors showed frequency-dependent sign and magnitude, and nearly all were significant (p_perm_ < 0.05 and parametric p < 0.05). All predictors failing to reach significance were observed for MEG-FC models including: myelin for gamma_hi_, caliber for alpha, and length for theta and beta. ß_myelin_ was negative for BOLD, delta, beta and gamma_lo_, in line with the global correlation of myelin and BOLD-FC(Nelson et al., 2023). ß_myelin_ was positive and higher in magnitude for theta and alpha. A similar sign flip was observed for caliber (ß > 0 for BOLD and most MEG bands; ß < 0 for gamma) and length (ß > 0 for BOLD and ß < 0 for most MEG bands).

The interaction terms tended to be the lowest ranked predictors across models, reflecting their general role of fine tuning the larger effects of their respective lower order terms.

Nonetheless, most interaction terms were significant and showed heterogeneous sign and magnitude across FC modality and frequency. Interestingly, the interaction term between myelin and length ranked 1^st^ in predictor dominance for alpha-band MEG and showed a large magnitude positive ß-weight indicating that the role of myelin is strengthened for longer edges. We expand on these interactions with myelin in subsequent sections.

These results communicate two important points about the structure-function relationship of the brain. First, white matter myelin is an important predictor of whole-brain FC showing comparable predictive power to the much more extensively studied white matter edge features caliber and length. Second, the specific roles of white matter features including myelin in mediating brain function depend on the cellular processes contributing to the measurement technique (haemodynamic vs electromagnetic) and the timescale of the signal (frequency bands).

### Haemodynamic and electromagnetic connectivity show distinct, spatially varying relationships with white matter myelin

Several reports have demonstrated regional heterogeneity in SFC using conventional SC directly or analytic measures derived therefrom(Baum et al., 2020; Liu et al., 2023; Preti & Van De Ville, 2019; Vázquez-Rodríguez et al., 2019; Zamani Esfahlani et al., 2022). Given our observations above and the diversity of mechanisms by which myelin supports brain function, we next sought to describe how the coupling of edge myelin and FC varies across brain regions. We ran the same multi-linear model at two additional spatial scales: (1) within pairwise combinations of the seven canonical resting-state networks(Thomas Yeo et al., 2011), and (2) within the connections of each individual node *i* to the remaining *j* ≠ *i* nodes. The limbic-somatomotor network pair was excluded for low degrees of freedom (d.f. < 10). We observed high correspondence between the network (**Figure 2**, **4**, **S1**, and **S2**) and node (**Figure 3** and **S3**-**S5**) levels. We thus focus this description on results from the network-level models as they are more robust (µ d.f. ≈ 875 vs 85) and efficient (n = 28 models vs 400).

**Figure 2.**
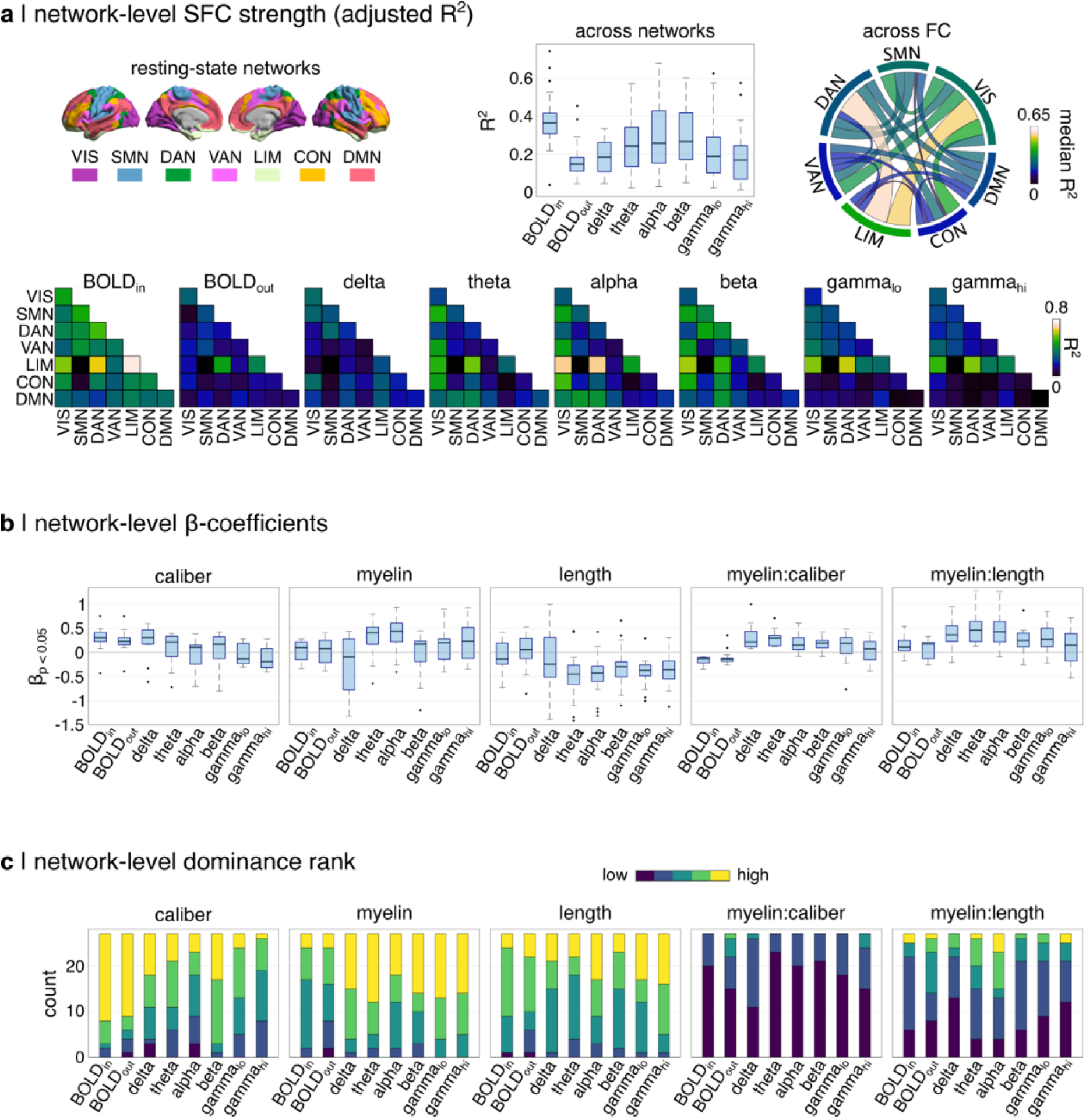
Relating white matter structural features to haemodynamic and electromagnetic connectivity at the level of pairwise combinations of resting-state networks.**(a)** SFC strength for pairwise combinations of resting-state networks defined using the Yeo 7-network atlas (top left). Boxplots (top middle) show the distribution of SFC across network pairs using standard elements: median line, 1^st^ and 3^rd^ quartile box limits, 1.5 interquartile range whiskers, and outlier data points. The circle plot (top right) shows the median value across FC networks with the network bars along the perimeter indicating the within-network values. SFC for every network pair and FC network is also shown in matrix form (bottom). **(b)** The distribution of standardized ß-coefficients and **(c)** the count of dominance rankings is shown for all predictors.

**Figure 3.**
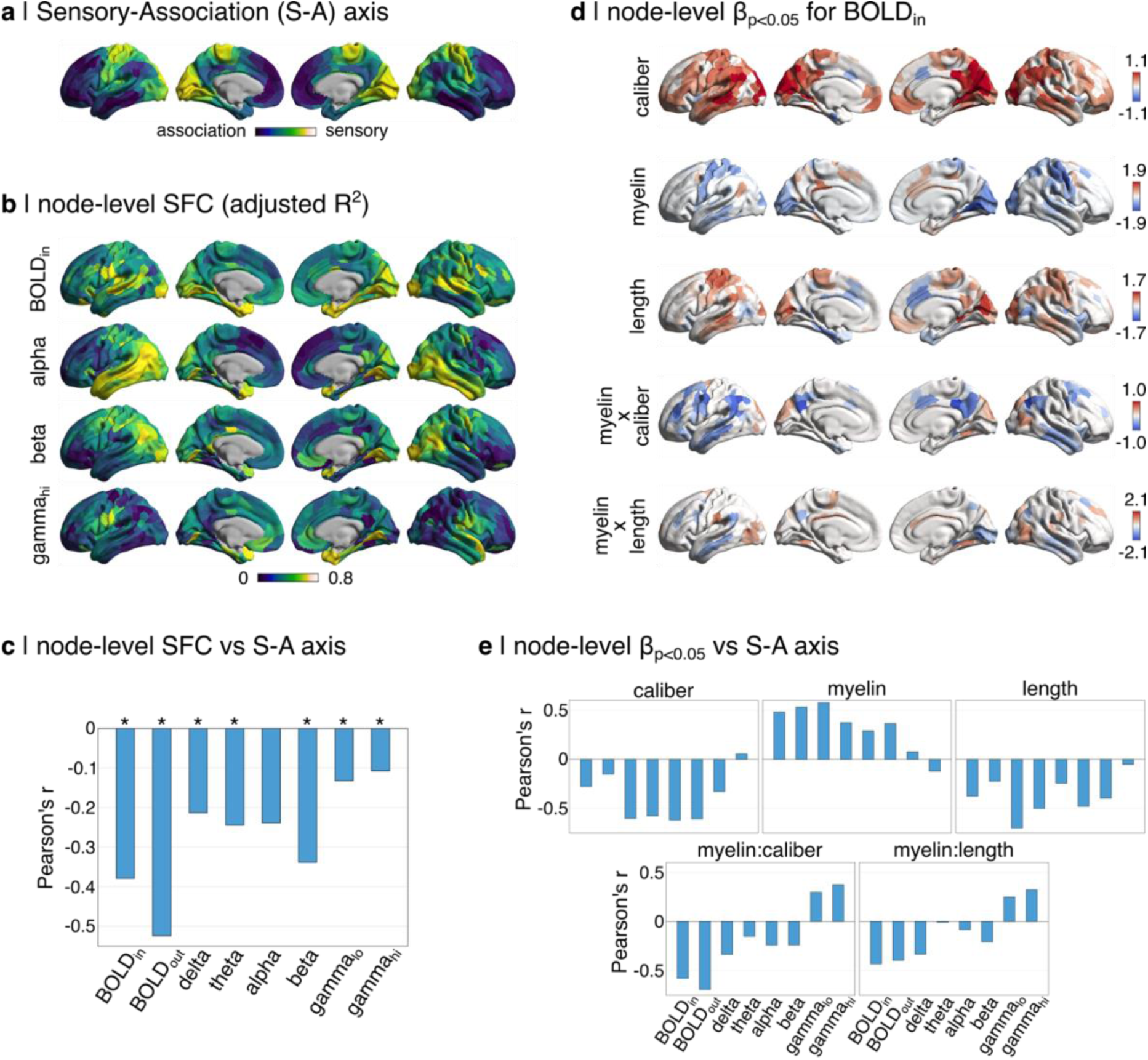
Alignment of node-level model coefficients with the sensory-association (S-A) axis. **(a)** The S-A axis is shown in the Schaefer-400 parcellation. Node values are shown for **(b)** SFC for several FC models, as well as the **(d)** ß-coefficients from the BOLD_in_ model for all structural predictors. Pearson’s correlation was used to quantify alignment for (**c**) SFC and (**e**) ß-coefficients for all predictors and FC models. All ß-values were filtered by parametric t-tests (p < 0.05). Asterisks additionally indicate significance from spatially-constrained permutation testing (p_perm_ < 0.05).

**Figure 4.**
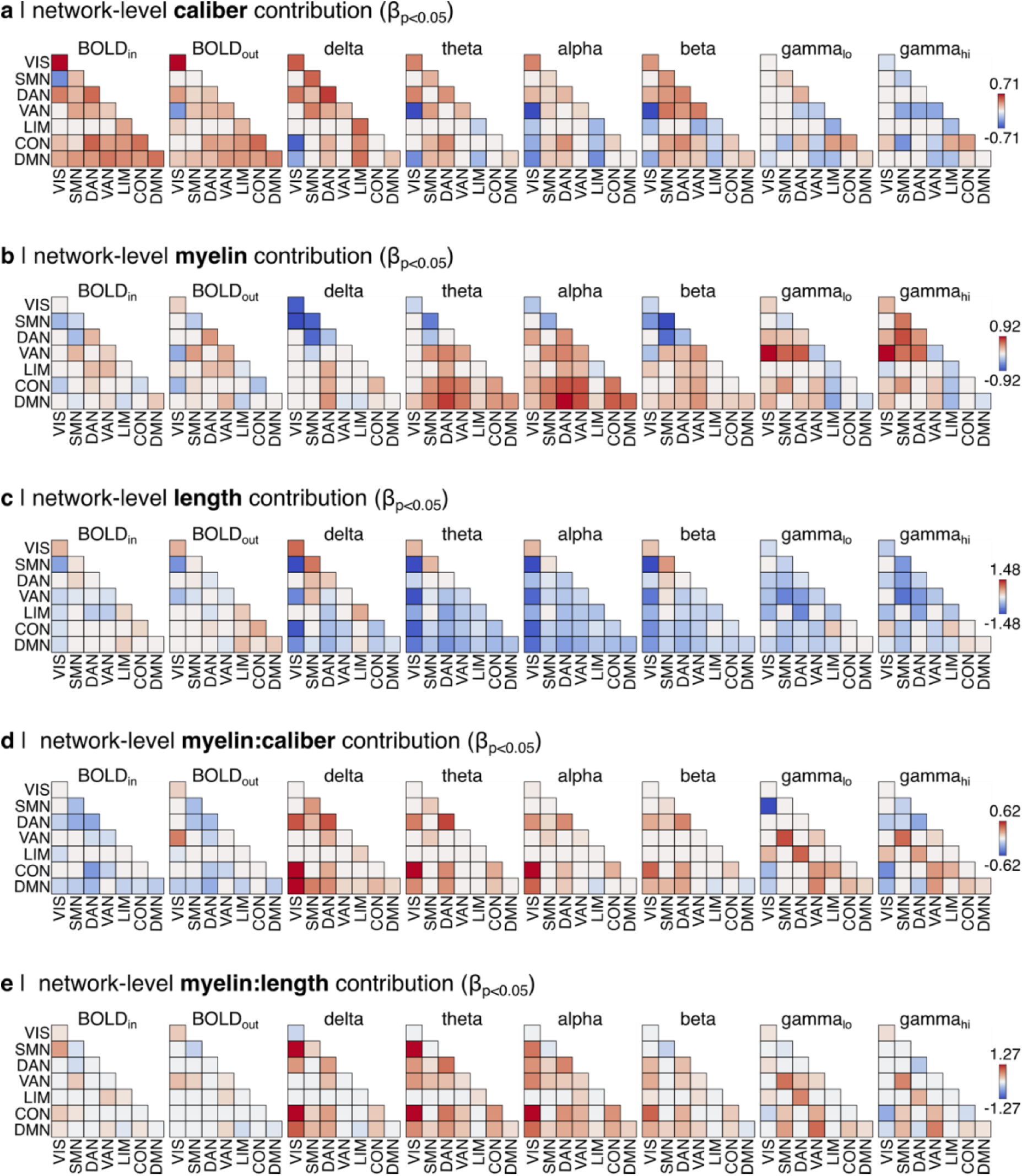
Standardized ß-coefficients for edge **(a)** caliber, **(b)** myelin, **(c)** length, (**d**) the interaction of myelin and caliber, and (**e**) the interaction of myelin and length at the pairwise network level for all FC models. Only significant values are shown (t-test p < 0.05).

Across network pairs and FC timescales, there was considerable heterogeneity in SFC (**Figure 2a**), as well as in the standardized ß-coefficients (**Figure 2b** and **4**) and predictor dominance rankings (**Figure 2c** and **S1**) for caliber, myelin, length, and both myelin interaction terms. SFC was strongest for BOLD_in_ followed by alpha and beta. Network pairings involving primary sensory regions (visual and somatomotor) showed stronger SFC than those involving association regions (default mode, control, and attentional). The strongest SFC was observed for the limbic-dorsal attention, limbic-visual, and limbic-limbic network pairs. At the node-level, SFC showed significant inverse correlations with the sensory-association axis for all FC networks, except alpha (**Figure 3c**). BOLD_in_ showed the most spatially distributed pattern of strong SFC, with the highest values spread across central, parietal, temporal, and occipital cortical regions (**Figure 3b** and **S3**).

The standardized ß-coefficients for caliber, myelin, and length (**Figure 2b** and **4**) showed both positive and negative values across network pairs and an interesting frequency dependence. ß_caliber_ showed a gradual transition from mostly positive to negative values when moving from low-to-high frequency FC (including BOLD), while ß_myelin_ showed an opposite trend: balanced for BOLD and delta, and more positive for theta through gamma. ß_length_ showed predominantly negative values, with exceptions for BOLD-FC and unimodal networks for most models.

Among network pairs, the spatial pattern of positive and negative ß_myelin_ was on average inversely aligned with the canonical unimodal-transmodal gradient of cortical organization: positive for network pairs involving transmodal association networks and negative for unimodal (**Figure S2a**). In contrast, ß_caliber_ was on average positive within a given network and mixed between networks regardless of their position on the gradient. Interestingly, the ß-coefficients for both myelin and caliber deviated from these average trends for gamma: ß_myelin_ was mostly positive including for unimodal networks, and ß_caliber_ was mostly negative including within-network (**Figure 4a** and **4b**). The ß-coefficients for both interaction terms were significant across most network pairs (**Figure 4d** and **4e**). ß_myelin:caliber_ was mostly negative for BOLD and positive for MEG. ß_myelin:length_ was mostly positive and tended to be larger. These interactions are explored in more detail in the subsequent section.

At the node-level, ß_caliber_ and ß_length_ were inversely correlated and ß_myelin_ positively correlated with the S-A axis across FC networks (**Figure 3e**). These trends were weakest or absent in the highest frequency gamma. ß_myelin_ tended to be larger in magnitude than ß_caliber_ (**Figure S5b**).

We also observed a frequency-dependence for the network-level predictor dominance rankings of caliber, myelin, length, and both myelin interaction terms (**Figure 2c** and **S1**).

Caliber tended to rank 1^st^ and myelin 2^nd^ for BOLD. As FC frequency increased, length became more dominant and caliber became less dominant, such that myelin and length shared dominance for most network pairs in alpha and gamma. Beta resembled BOLD but with greater influence of length. Myelin showed strong interactions, especially with length for alpha and theta. At the node-level, caliber tended to dominate in frontal and temporal regions and myelin in central, parietal and occipital regions for BOLD (**Figure S4**).

With these results, we have revealed distinct spatial patterns of structure-function coupling for each frequency band. Furthermore, we have demonstrated how these relationships can be decomposed to examine the influence of individual predictors (white matter features). Despite the heterogeneity of spatial patterns, there is generally a shared alignment with the sensory-association axis across predictors and most frequencies of FC.

### Antagonistic gradients of structure-function coupling and white matter myelin

Next, we dove deeper into the strong interactions with myelin reported above by asking how the relationships of caliber and length with FC may be different for high vs low myelin edges. We partitioned network edges into 5 equal-sized bins (n ≈ 3667) according to the magnitude of their myelin weight (**Figure 5a**). Within these bins, we computed Pearson’s correlations of the z-scored SC and FC edge weights, as well as a version of the previously described multi-linear model excluding the myelin-interaction terms.

**Figure 5.**
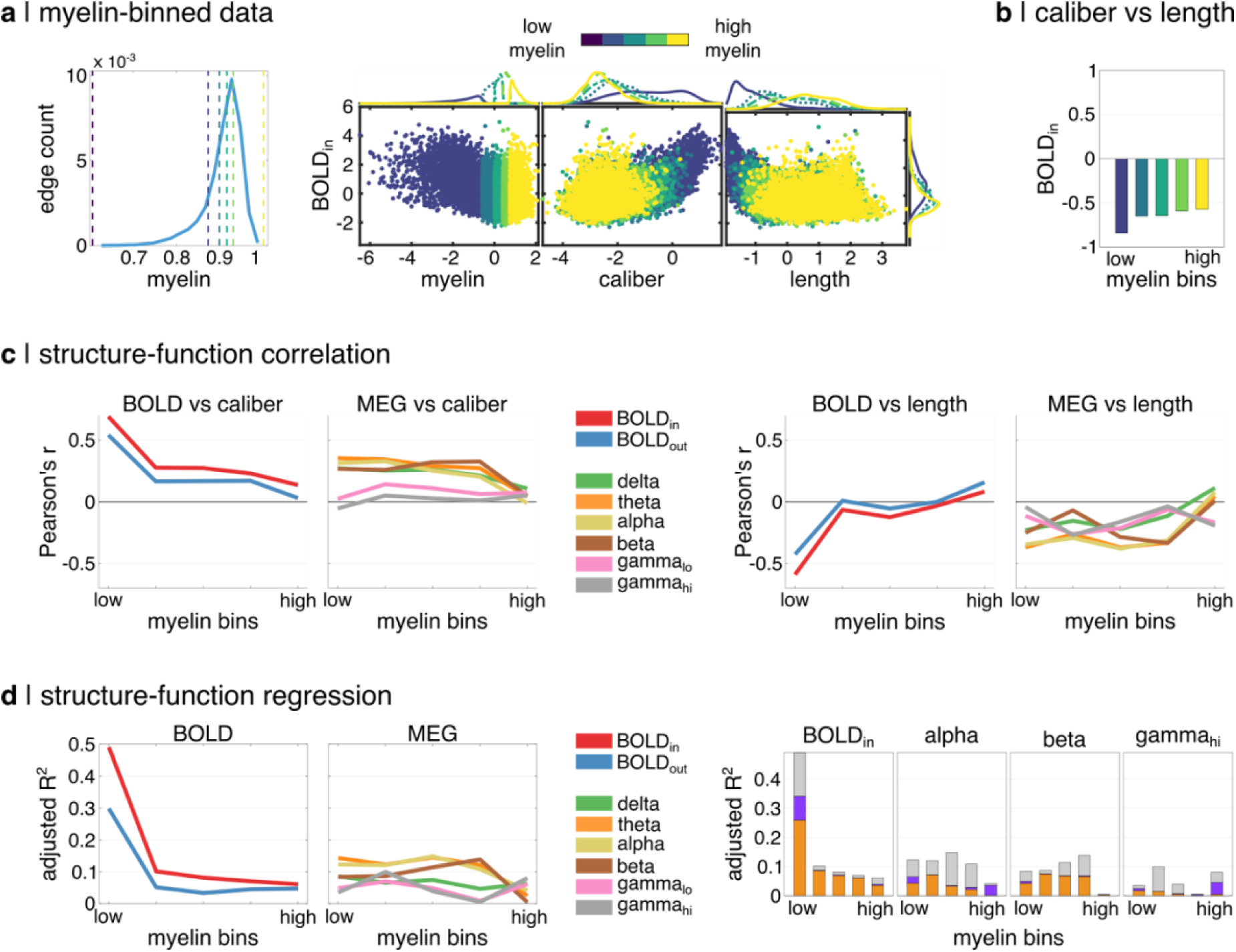
The coupling of white matter structure and FC weakens along the gradient of white matter myelin. **(a)** Network edges were partitioned into 5 equal-sized bins by edge myelin magnitude. Each bin contained 20% or ≈ 3667 network edges. **(b)** The Pearson correlation of caliber and length across myelin bins. **(c)** The Pearson correlation of caliber (left) and length (right) with both BOLD-FC and MEG-FC is shown across myelin bins. **(d)** SFC (adjusted R^2^) across myelin bins is shown for all FC networks (left) and for a subset including predictor dominance ratios (right).

While the relationship between caliber and length remained relatively constant with increasing myelin magnitude (**Figure 5b**), both length and caliber uncoupled from BOLD-FC and low-intermediate frequency MEG-FC (delta, theta, alpha, beta) (**Figure 5c**). This trend is in line with a potential role for white matter myelin in mediating the relationship between FC and white matter edge macrostructure. For BOLD-FC, this uncoupling effect was gradual for caliber, whereas the correlation of length and BOLD-FC was already near 0 in the 2^nd^ myelin bin. For low-to-intermediate frequency MEG, uncoupling of either length or caliber did not occur until the final bin with the highest myelin magnitude. The gamma bands remained relatively uncoupled from caliber independent of myelin and showed distinct patterns of coupling with length.

For BOLD, SFC was much stronger when myelin was lowest (BOLD_in_ R^2^ = 0.49; BOLD_out_ R^2^ = 0.30) and decreased dramatically with increasing myelin magnitude (**Figure 5d**). The predictor contributions show that this reduction in SFC is driven by a large reduction in the impact of caliber and a complete elimination of length. For theta, alpha, and beta bands, caliber and length continue to drive consistent SFC until a drop off in the final high myelin bin. Delta and gamma deviate from this trend by showing weak relationships with structure in all myelin bins. These results provide evidence for a progressive dissociation of white matter structural and functional connectivity with increasing white matter myelin content. The modality-specificity of this effect highlights the multi-layered relationship between brain structure and function.

### A multi-feature structural model predicts robust general FC patterns but weak subject-specificity

Our final goal was to assess the generalizability of the full linear model with interaction terms to unseen subject-level data from BOLD_in_ (n = 10; 5 subjects, 2 scans, inter-scan interval < 3 weeks). In a leave-one-out analysis by test subject (*S_x_*), group-level SC and FC networks were computed excluding data from both scans, and multi-linear modeling was performed at the pairwise network level. The resultant group-level ß-coefficients were then combined with the unseen subject-level SC data to compute predicted FC, which was correlated with the corresponding empirical FC for both time points using Pearson’s correlation (r_true_ = *FC_predicted_(S_x_)* vs *FC_empirical_(S_x_)*). We assessed the significance of r_true_ using a 2-level approach. (1) The ability of the model to produce a general FC pattern was assessed by correlating the predicted FC for *S_x_* with permuted versions of their empirical FC (n = 1000; r_perm_ = *FC_predicted_(S_x_)* vs *FC_empirical-shuffled_(S_x_)*). Shuffling was performed within network pairs to preserve spatial relationships. (2) The ability of the model to produce subject-specific FC patterns was assessed by correlating the predicted FC for *S_x_* with the empirical FC for every other subject y ≠ x (r_spec_ = *FC_predicted_(S_x_)* vs *FC_empirical_(S_y_)*). For both tests, correlations were performed over all pooled edges (**Figure 6a** and **6b**) and within network pairs (**Figure 6c**). Significance was assessed using one-sided p-values and z-statistics.

**Figure 6.**
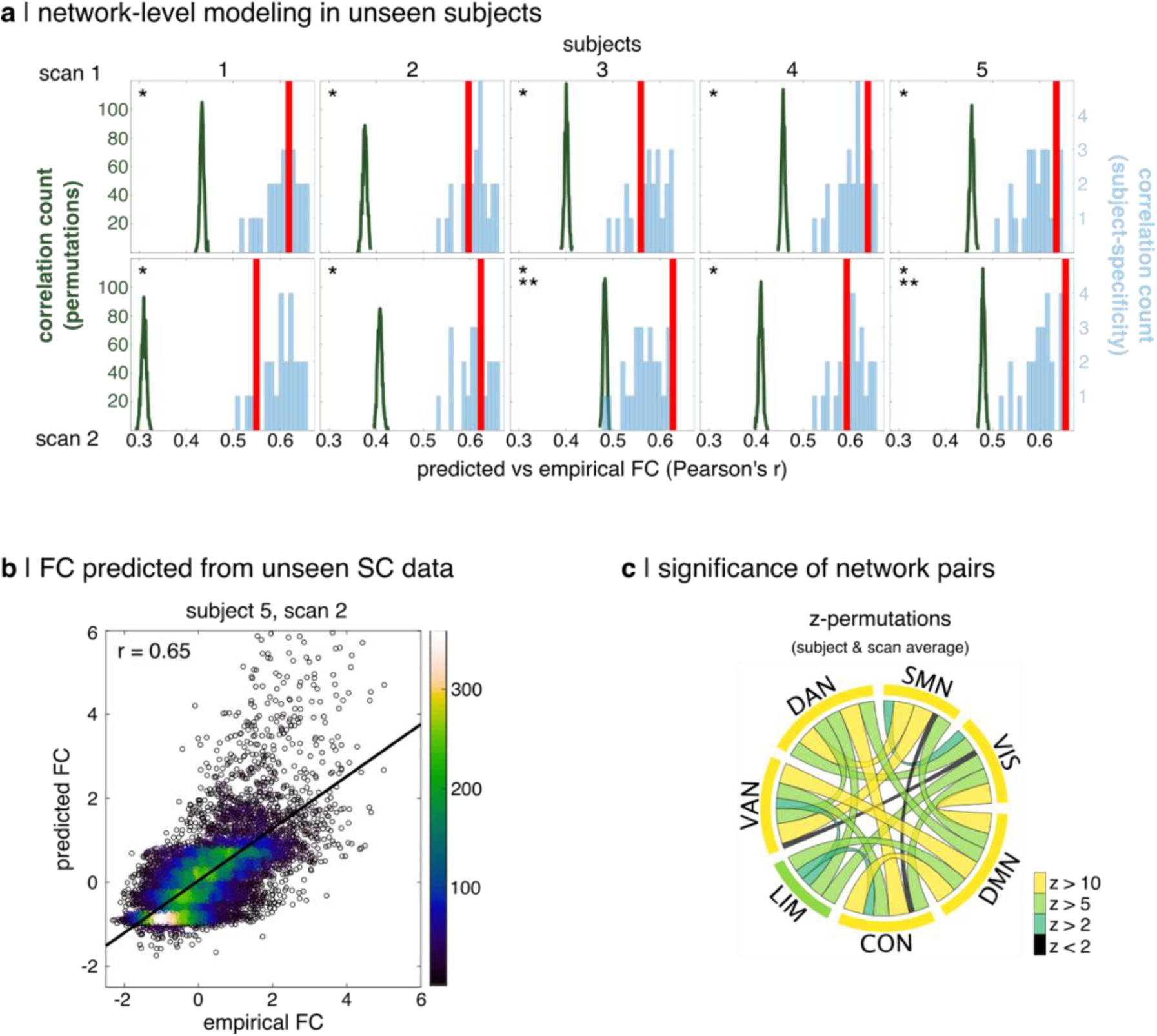
Generalizability and subject-specificity of the full regression model of BOLD_in_ at the network level. **(a)** True correlations of empirical and predicted FC (r_true_; red bar) are overlaid on the distributions of test correlations for generalizability (r_perm_; forest green) and subject-specificity (r_spec_; light blue). Single (*) and double asterisks (**) indicate p_perm_ and p_spec_ < 0.05, respectively. **(b)** Empirical vs predicted FC for the highest true correlation (r_true_ = 0.65). **(c)** Generalizability significance for each network pair was computed as z_perm_ = (r_true_ – mean_r_perm_) / std_r_perm_. No network pair showed significant subject-specificity.

For every subject and scan, r_true_ was significantly higher than r_perm_ (p_perm_ = 0.001), indicating that the pattern of predicted FC was more similar to empirical FC than random sequences of FC edges controlling for spatial relationships. In contrast, r_true_ was only significantly higher than r_spec_ for 2 of 10 tests (p_spec_ = 0.001, 0.041). When averaging across subjects and scans, r_true_ was significantly higher than r_perm_ (z > 2) for nearly every network pair and was an extreme outlier (z > 10) for 12 of 29 pairs which included all except the limbic network. No network pairs showed significantly higher r_true_ than r_spec_ (all z < 1). Thus, the predicted FC from our multi-linear structural model of function showed strong general patterns of FC but weak subject-specificity.

### Sensitivity analyses

Finally, we test the extent to which our results may be sensitive to our specific dataset or processing choices. Our main results showed good consistency when replicated using an independent multi-modal MRI dataset(Royer et al., 2022) to define the structural features and the BOLD_in_ network (**Figure S8**). The edge myelin connectome was quantified using the same method (tractometry) but a different metric in these data (R_1_ = 1/T_1_; longitudinal relaxation rate). We also observed consistency using the lower-resolution Schaefer-200 parcellation (**Figure S6**). All processing steps were identical, except a more conservative threshold was used for consistency-based filtering. In our work, the COMMIT framework was used to derive an estimate of edge caliber, which can be interpreted as quantifying the strength of white matter connectivity. However, the number of streamlines output from streamline tractography is more often used to quantify SC in structure-function models. To support comparability to previous literature, we additionally show that our results are consistent when using the number of streamlines as an estimate of caliber (**Figure S7)**.

## Discussion

The white matter tracts connecting gray matter neuronal populations play a central role in supporting the synchronous neural activity necessary for healthy brain function(Sonya Bells et al., 2017; Filley & Fields, 2016). Yet, we still know very little about the mechanisms mediating this relationship, in part, due to a lack of requisite biological detail in structure-function brain models(Suárez et al., 2020). Here, we provide the most thorough account to date of the relationship between multi-feature white matter structural connectivity and multi-modal functional connectivity. We find that white matter myelin is strongly predictive of FC with coupling patterns that vary regionally and across timescales of neural function.

Mounting evidence supports a role for white matter myelin in mediating macroscale brain function, which complements the established relationship between intracortical myelin and BOLD-FC(Huntenburg et al., 2017). White matter myelin has been used to predict latent components of MEG-FC(Messaritaki et al., 2021). Changes in white matter myelination have been linked causally with disruptions in network connectivity in mouse models(Asleh et al., 2020; Saskia Hübner et al., 2017) and with working memory impairment using computational modeling(Ibañez et al., 2023). Using MRI-based white matter myelin markers, changes in both network myelination(Metzler-Baddeley et al., 2017) and myelin-weighted network topology(Caeyenberghs et al., 2016) have been detected following working memory training.

Connectivity networks weighted by white matter myelin have also been shown to differ markedly in topology and correlation with FC from conventional SC networks supporting their complimentary relationships with function(Boshkovski et al., 2021; Nelson et al., 2023). We extend this work by describing in detail how white matter myelin shapes the relationship between structural and functional human brain networks.

### White matter myelin as a modulator of structure-function coupling

Structure-function coupling strength was inversely related to the sensory-association axis, in line with previous studies which did not include white matter myelin(Baum et al., 2020; Liu et al., 2023; Osmanlıoğlu et al., 2019; Preti & Van De Ville, 2019; Vázquez-Rodríguez et al., 2019; Wang et al., 2019; Zamani Esfahlani et al., 2022). The oft-observed uncoupling of white matter SC from FC along this gradient is thought to be related to a progressive untethering of the cortex from underlying molecular gradients throughout evolution, which has occurred to the greatest extent in association cortex in humans(Buckner & Krienen, 2013). This –– along with a similar pattern in development(Hill et al., 2010) –– has been linked to cortical features including myelin(Glasser et al., 2014; Glasser & Van Essen, 2011), cell morphology(Elston, 2003; Glasser & Van Essen, 2011), and transcriptomic alignment(Burt et al., 2018), as well as the genetic heritability of FC(Giles L. Colclough et al., 2017). But what is the role of white matter itself in the uncoupling of SC and FC?

Myelin showed the opposite relationship with BOLD and MEG FC across the sensory-association axis relative to caliber and length. We observed consistently strong interactions with white matter myelin for both caliber and length. When stratifying by myelin content, BOLD structure-function coupling was much stronger when myelin was lowest. Both length and caliber uncoupled from BOLD-FC with increasing myelin, while remaining relatively strongly correlated with each other. These results suggest a role for white matter myelin as a *moderator* of structure-function coupling. When myelin is low, the neural synchrony quantified by FC may be more heavily influenced (i.e., constrained) by other physical properties of white matter connections, such as their length and caliber. Increasing myelination may gradually compensate for variability in these other features, such that a sufficient threshold of myelin has the net effect of equalizing functional synchrony across a diverse range of tract profiles: i.e., a flattening of the structure-function gradient. In other words, white matter myelin may confer functional flexibility or resilience(Noori et al., 2020; Park & Lefebvre, 2020), whether it be to normal or pathological variability in white matter structural properties. Future work may explore this further, e.g., in aging(Slater et al., 2019; Yeatman et al., 2014). Thus, we provide evidence for a modulatory role for white matter myelin in the progressive decoupling of function from its underlying white matter macrostructural constraints, which may contribute to the greater flexibility of brain function characteristic of association networks.

Previous work has established that the fMRI signal is best described by a spatially varying superposition of multiple MEG frequency bands(Shafiei et al., 2022; Tewarie et al., 2016). The modulatory effect of myelin on structure-function coupling was consistent in the out-of-sample BOLD data but was heterogeneous across MEG frequency bands: mostly absent from the gamma bands, and mostly present for low-intermediate frequency bands, although only for edges with the highest myelin content. Taken together, these results suggest that FC for low-to-intermediate frequency MEG may be more strongly constrained by white matter macrostructure, as decoupling is only observed at a higher myelin threshold. Furthermore, FC for high-frequency MEG may be fundamentally decoupled from white matter caliber, which is additionally supported by the relative unimportance of caliber in the prediction of gamma-FC. This is consistent with a more local and transient nature, which is more strongly dependent on cortical – – as opposed to white matter ––structural features(Buzsáki & Wang, 2012). Thus, the stronger modulatory effect observed for BOLD-FC may reflect the filtered integration of heterogeneous modulation of structure-function coupling by white matter myelin across multiple neurophysiological rhythms of brain function.

In sum, we provide robust evidence that white matter myelin has a frequency-dependent modulatory effect on structure-function coupling, which may contribute to the gradual uncoupling of white matter structure and FC along the sensory-association gradient.

### Differential roles for white matter myelin in supporting BOLD-FC across the unimodal-transmodal gradient

The FC of a given node pair derived from BOLD-fMRI is an indirect measure of synchronized neural activity, which will intuitively depend on signal transmission speed and distance. Myelin was a stronger predictor of BOLD-FC than length both globally and for most network pairs. On face value, this would suggest that white matter myelin would be a useful proxy for *in vivo* conduction delays; however, previous work has provided evidence to the contrary(Mancini et al., 2021).

Myelin generally showed a balanced relationship with BOLD-FC: mostly negative in unimodal and positive in heteromodal networks. This implies that the strength of neural coupling measured with BOLD-fMRI is associated with lower tract myelination in unimodal and higher myelination in heteromodal networks, which agrees with previous work(Boshkovski et al., 2021; Sydnor et al., 2021). Whereas the hub regions(Martijn P. van den Heuvel & Sporns, 2013) driving the inter-modular integration characteristic of heteromodal networks tend to be connected by long-range white matter tracts, the more segregated processing attributed to unimodal networks tends to be supported by dense local circuits with shorter connections(Betzel & Bassett, 2018; Sepulcre et al., 2010; Sydnor et al., 2021). In this light, our results could indicate that BOLD-FC in heteromodal networks is more reliant on high myelination of white matter connections for temporal coordination between distant brain regions.

Alternatively, higher myelination may help reduce the energetic costs of topologically central long-range connections(Bullmore & Sporns, 2012; Harris & Attwell, 2012). A related but separate consideration is adequate trophic support, especially when viewing the insulating effect of myelin from the perspective of metabolic isolation(Nave, 2010). Indeed, mounting evidence suggests that oligodendrocytes directly support axonal energy metabolism independent of their role in modulating conduction velocity(Asadollahi et al., 2024; Fünfschilling et al., 2012; Y. Lee et al., 2012; Narine & Colognato, 2022; Nave et al., 2023; Nave & Werner, 2014; Ramos-Cabrer et al., 2025; Saab et al., 2016). Thus –– alongside temporal coordination –– both energetics and trophic support may contribute to the observed differential association between myelination and BOLD-FC across the S-A axis.

### Connecting white matter and neural synchrony

Our approach provides the unique opportunity to speculate on the interplay between white matter structure and neural synchrony across functional timescales. MEG captures oscillatory neural activity at higher frequencies than BOLD-fMRI and should thus be more sensitive to increasing conduction delays in longer white matter tracts. Indeed, we observed that length increased in importance shifting from BOLD to MEG to become the dominant predictor overall across MEG frequency bands. Myelin also increased in importance and showed predominantly positive associations in the highest frequency FC. This could be argued to bolster the support of conduction velocity, or perhaps to reflect increasing energy demands at higher frequencies(Lord et al., 2013; Saab et al., 2016; Trevisiol et al., 2017). In line with past work(Hagmann et al., 2008; Koch et al., 2002), caliber dominated the prediction of BOLD-FC showing almost exclusively positive associations. However, as the frequency of brain function increased, caliber decreased in predictive power and shifted to predominantly negative associations with gamma-FC. Taken together, these observations support an intuitive shift in priorities across functional timescales: from higher bandwidth at lower frequencies to greater temporal precision and/or metabolic demands at higher frequencies.

Myelin and length shared dominance in predicting alpha and gamma across network pairs. The association of myelin and FC was inverted between these frequency bands as well: mostly positive in unimodal and negative in heteromodal for gamma_hi,_ and the inverse for alpha. Gamma is associated with local, bottom-up sensory processing and alpha with top-down modulation over longer distances(Baillet, 2017; Fries, 2015; Scheeringa & Fries, 2019). Thus, for intermediate-frequency alpha, the dual-dependence of strong FC on both myelin and length may reflect the role of supporting synchronization over longer distances –– as in low-frequency BOLD-FC –– alongside a greater reliance on high temporal fidelity. Whereas for high-frequency gamma, the strong dependence of synchrony on physical proximity is evident in the dominance of length, and the significant contribution of myelin reinforces its importance for both timing and metabolic support.

In sum, our results support a frequency-dependent relationship between white matter structure and neural synchrony. In the lowest-frequency BOLD-FC, a global emphasis on bandwidth is suggested by a strong reliance on caliber, and high myelination could be argued to promote temporal coordination and metabolic support for topologically central long-range connections. As the frequency of FC increases, the global reliance on bandwidth is gradually reduced and eventually overcome by a greater dependence on temporal precision, which is supported by the increasing dominance of length and myelin.

### Advantages of a multi-feature, network-level structure-function model

In line with past work(Liu et al., 2023; Zamani Esfahlani et al., 2022), structure-function coupling strength and the correlation of empirical with predicted FC increased when moving from a single global model to smaller regional models. We found the strongest structure-function coupling in limbic network pairs, however the limbic network is typically among the lowest ranked. Previous studies have defined network-level coupling by aggregating over node-level models(Liu et al., 2023; Vázquez-Rodríguez et al., 2019; Zamani Esfahlani et al., 2022), but each node-level model contains edges from multiple network pairs. In contrast, our network-level model focused specifically on edges between pairwise combinations of resting-state networks, which reduced the overall number of models by an order of magnitude and increased the number of data points in each model. Thus, our network-centric approach was more robust, efficient and yielded a more nuanced perspective of structure-function coupling. A complementary but uniquely informative example can be seen in the somatomotor network. Past work has reported strong average structure-function coupling here(Liu et al., 2023; Zamani Esfahlani et al., 2022), and our results generally support this, however with a caveat. The somatomotor network showed no coupling with the limbic and very low coupling strength with control and default mode networks. Therefore, relative to the ventral attention network, which is typically ranked among the weakest in structure-function coupling, the somatomotor network had lower total coupling strength across all network pairs.

Our analysis also deviated from past work by including multiple white matter structural features and their interactions. Structure-function coupling in BOLD was strongest for network pairs including the visual and dorsal attention networks. The finding in the visual network is corroborated by multiple investigations(Liu et al., 2023; Vázquez-Rodríguez et al., 2019; Zamani Esfahlani et al., 2022), however, the dorsal attention network does not typically rank highly in terms of structure-function coupling strength among resting-state networks(Vázquez-Rodríguez et al., 2019). While most network pairs with the dorsal attention network showed edge caliber as the dominant predictor, large magnitude ß-coefficients were observed for both myelin and length for the network pair with the strongest coupling overall: dorsal attention and limbic. This suggests that the strong structure-function coupling observed with the dorsal attention network in our study may result from our multi-feature approach which included myelin. Another example can be provided using our beta-FC results. While BOLD-FC is best characterized by a mixture of MEG frequency bands, the beta band is thought to be a primary driver of their correspondence(Shafiei et al., 2022). We observed a similar balanced association between myelin and both BOLD and beta FC which showed roughly the same distribution across resting-state networks. In contrast, caliber, length, and both interaction terms showed noticeable differences, suggesting that the strong correspondence between FC in BOLD and beta may be related to overlap in their relationships with white matter myelin. Future work could further explore this possibility using EEG, which is more widely available and also related to SC(Babaeeghazvini et al., 2021).

In sum, the pairwise network-level of analysis yielded a nuanced perspective of regional structure-function coupling while maximizing model robustness and efficiency, and our multi-feature account of white matter structure facilitated deeper interpretation of the relationship between brain structure and function.

### Limitations

The principal limitation of this work was the exclusion of some parts of the network from our analysis. (1) Network edges with SC = 0 were excluded, which constitutes more than 80% of the FC network. This is unavoidable when modeling edgewise relationships with structural edge weights directly, as structural brain networks are inherently sparse. This could be addressed in future work using communication modelling(Seguin et al., 2020, 2023). (2) The subcortex and cerebellum were excluded, as these are associated with increased artifact and a lack of standardization for their inclusion in whole brain analyses. (3) Cortical features were excluded in order to focus our analysis on the white matter. Future extensions of this work could integrate these other network components.

A further limitation of the present work is that the MEG FC data were derived from a separate cohort than the structural connectivity used in the modeling. However, all analyses were conducted at the group level, and our interpretations rely on broad population-level trends rather than subject-specific structure–function coupling. We made no inference at the level of individual tracts or anatomical features, and no subject-level predictions were performed using the MEG data. As such, the use of out-of-sample MEG data is unlikely to materially affect the central conclusions. Future work should integrate within-subject multimodal datasets that jointly acquire structural, fMRI, and MEG/EEG measurements. Such data would enable subject-level analyses of myelin-sensitive communication, allow stronger inferences about individual variability in structure–function coupling, and facilitate testing whether the group-level principles identified here also hold at the level of single participants. In addition, acquiring myelin-sensitive structural measures and electrophysiological activity within the same individuals would enable this framework to be extended to clinical populations with myelin alterations (e.g., demyelinating, neurodevelopmental, or neurodegenerative conditions). Such datasets would make it possible to assess how individual variability in myelin integrity relates to large-scale functional coherence or synchrony, and whether structural disruptions associated with altered myelination are reflected in corresponding changes in electrophysiological coupling.

Another limitation was the use of streamline-tractography and tractometry to compute SC networks. This imparts well known biases(Maier-Hein et al., 2017; Nelson et al., 2023; Schilling et al., 2019). We attempted to mitigate this to the fullest extent possible through careful filtering at both the streamline and connectome levels. We also used a group-consensus approach. This improves the stability of the analysis and focuses on population-level trends at the expense of subject-specificity, e.g., through personalized parcellations(Kong et al., 2021).

Our investigation utilized a single myelin metric derived from MTsat, which provides an estimate of myelin density. Future work could expand on this by measuring myelin with alternate metrics such as the tract g-ratio(Lu et al., 2025; Mancini et al., 2018; Stikov et al., 2015; West et al., 2016). This would provide a complimentary perspective of white matter myelin and further illuminate its coupling with FC. Similarly, future work could expand on the method for estimating caliber e.g., axonal caliber(Barakovic et al., 2021).

Throughout, we have interpreted the sign and magnitude of standardized ß-coefficients in terms of the relationship between predictors (white matter features) and the dependent variable (FC). We recognize that this must be done cautiously, as the ß-coefficients from a given model are sensitive to the predictors included and their collinearity, which is unavoidable in our context. To account for this, all variables were normalized and centered prior to modeling, and dominance analysis was used to quantify relative predictor contributions independent of collinearity.

## Conclusion

This work constitutes the most complete description of the relationship between white matter structural features –– including myelin –– and functional connectivity across time scales and brain regions. We detail the complex interplay between white matter structure and neural synchrony and describe how it evolves across the neurophysiological rhythms of the brain.

Crucially, we provide robust evidence of a frequency-dependent modulatory role for white matter myelin on structure-function coupling along the sensory-association gradient. In all, our contributions highlight the necessity of a multi-feature account of white matter which includes myelin for investigations of the structure-function relationship of the brain.

## Materials and Methods

### Data acquisition

This study was approved by the Research Ethics Board of McGill University, and all participants provided written informed consent. Multi-modal MRI data were collected in a cohort of healthy volunteers (n = 30; 14 men; 29±6 years of age) on a 3 tesla Siemens Magnetom Prisma-Fit scanner equipped with a 64-channel head coil. All sequences used the Siemens *Prescan*

*Normalize* option to generate maps with minimal receive B1-field bias. The protocol was as follows:

- T_1_-weighted (T_1_w) anatomical: 3D magnetization-prepared rapid gradient-echo sequence (MP-RAGE; 1.0 mm isotropic; TE/TI/TR = 2.98/900/2300 ms; FA = 9°; iPAT = 2).
- T_1_ relaxometry: sparse 3D-MP2RAGE(Marques et al., 2010; Mussard et al., 2020) sequence (1.0 mm isotropic; TE/TI_1_/TI_2_/TR = 2.66/940/2830/5000 ms; FA_1_/FA_2_ = 4°/5°; TF = 175; undersampling factor = 4.6.
- Resting-state fMRI: multiband accelerated 2D-BOLD gradient echo echo-planar sequence (2.6 mm isotropic; TE/TR = 30/746 ms; MB factor = 6; FA = 50°). Two spin-echo images with AP and PA phase encoding were additionally acquired for distortion correction (2.6 mm isotropic; TE/TR = 60.6/6160 ms; FA = 90°).
- Magnetization transfer-weighted 3D gradient-echo sequence (1.0 mm isotropic; TE/TR = 2.76/27 ms; FA = 6°; iPAT = 2; partial Fourier = 6/8; MT pulse parameters: 12-ms Gaussian pulse, 2kHz off-resonance, B1_rms_ = 3.26μT.
- Multi-shell diffusion-weighted imaging (DWI): 2D pulsed gradient spin-echo echo-planar imaging sequence composed of three shells with b-values 300, 1000 and 2000 s/mm^2^ and diffusion directions 10, 30, and 64, respectively (2.6 mm isotropic; TE/TR = 57/3000 ms; iPAT = 3; partial Fourier = 6/8). Two b = 0 images with AP and PA phase encoding were additionally acquired to facilitate distortion correction.
- A B ^+^ map (2.5 x 2.5 x 3mm^3^; TE/TR = 2.22/20k ms; FA = 8°) for ΔB_1_^+^ correction of MP2RAGE & MTsat maps.

### Data processing

A custom version of the multi-modal processing pipeline *micapipe* (v0.1.5)(Cruces et al., 2022) was used in the processing of all diffusion, anatomical, and functional images. This pipeline makes use of multiple software packages including: *AFNI*(Cox, 1996; Cox & Hyde, 1997), *FSL*(Jenkinson et al., 2012; S. M. Smith et al., 2004; Woolrich et al., 2009), *ANTs*(B. Avants et al., 2009; Brian B. Avants et al., 2014), *FreeSurfer*(Dale et al., 1999; Fischl, 2012; Fischl et al., 2002, 2004; Fischl, Sereno, & Dale, 1999; Fischl, Sereno, Tootell, et al., 1999), and *MRtrix3*(J. D. Tournier et al., 2012, 2019).

#### T_1_w anatomical MRI (MPRAGE)

T_1_w images were deobliqued, reoriented to standard neuroscience orientation (LPI), N4 bias field corrected(Tustison et al., 2010), intensity normalized, and skull stripped(Hoopes et al., 2022). Model-based segmentation of subcortical structures was performed with FSL FIRST(Patenaude et al., 2011) and tissue types were classified using FSL FAST(Zhang et al., 2001). A five-tissue-type image segmentation was generated for anatomically constrained tractography(R. E. Smith et al., 2012). Cortical surface segmentations were generated with the *FreeSurfer* (v7.2) *recon-all* pipeline with manual edits to remove dura mater.

#### Resting-state fMRI

After dropping the first 5 TRs from the main scan, all fMRI images (including field maps) were reoriented to standard neuroscience orientation (LPI) and motion corrected within-scan by registering all volumes to their respective temporal mean. Framewise displacement was computed for all main scan volumes, and a threshold (3^rd^ quartile + 1.5 * Interquartile Range) was used to identify volumes with large motion artifacts. The main-phase and reverse-phase field maps were used to correct the main fMRI scan for geometric distortions due to magnetic field inhomogeneity(Andersson et al., 2003; Graham et al., 2017). All subsequent processing involves only the main fMRI scan. Brain extraction(S. M. Smith, 2002) was performed, followed by registrations to native FreeSurfer space with *bbregister*(Greve & Fischl, 2009) and with the T_1_w image using a multi-stage method (rigid, affine, SyN) implemented in ANTs(B. B. Avants et al., 2008).

The fMRI data was denoised by applying a high-pass filter at a cutoff frequency of 0.01 Hz followed by Independent Component Analysis with FSL MELODIC(C.F. Beckmann & Smith, 2004; Christian F. Beckmann et al., 2005). All independent components (n = 3,825) were manually labeled by 2 independent raters, and components labeled as “noise” by both raters were removed (n = 2761). The denoised data was then used to calculate tissue-specific signals for gray matter, white matter, and cerebrospinal fluid tissue classes, before being mapped to native surface space and smoothed (Gaussian, FWHM = 6 mm). Linear regression was applied to the surface-based time series to remove nuisance signal resulting from motion, cerebrospinal fluid or white matter tissue.

#### Multi-shell DWI

The DWI data including field maps (b = 0 s/mm^2^ volumes with reverse phase encoding) were denoised(Cordero-Grande et al., 2019; Veraart, Fieremans, et al., 2016; Veraart, Novikov, et al., 2016), then corrected for Gibbs ringing(H. Lee et al., 2021). The field maps were then used to correct for susceptibility distortion, head motion, and eddy currents via FSL’s TOPUP(Andersson et al., 2003) and eddy(Andersson & Sotiropoulos, 2016). This was followed by B1^+^ bias-field correction(Tustison et al., 2010). The corrected DWI image was upsampled to match the T1w resolution (1 mm isotropic), and brain extraction was performed(S. M. Smith, 2002). These preprocessed data were used to estimate multi-shell and multi-tissue response functions(Christiaens et al., 2015; Dhollander et al., 2016, 2019), which informed the estimation of fiber orientation distributions (FOD) via spherical deconvolution(Jeurissen et al., 2014). All FODs were intensity normalized(Dhollander et al., 2021). A multi-stage (rigid, affine, SyN) registration(B. B. Avants et al., 2008) was computed with the T_1_w image to allow co-registration of anatomical images to DWI space.

#### T_1_ relaxometry and magnetization transfer saturation (MTsat) maps

A dictionary matching approach incorporating the B ^+^ map was applied to generate S_0_ and T_1_ maps from the MP2RAGE inversion 1, inversion 2 and UNI images. This produced ΔB_1_ ^+^ corrected S_0_ & T_1_ maps. An affine transformation (ANTs) was used to register the MP2RAGE denoised UNI image, the S_0_ and T_1_ maps, and the MT-weighted image to the T_1_w image. MTsat maps were computed using(Helms et al., 2008):

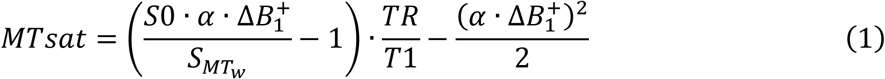

where 𝛼 is the excitation flip angle in the MT-weighted sequence, 𝑆_𝑀𝑇𝑤_ is the MT-weighted signal, and TR is the repetition time of the MT-weighted sequence. A gain factor of 2.5 was applied to the S_0_ maps to match the receiver gain of the MT-weighted gradient echo sequence for computing MTsat. ΔB ^+^ was corrected using a model-based approach(Rowley et al., 2021).

### Structural connectivity network reconstruction

#### Tractography and filtering

Anatomically constrained(R. E. Smith et al., 2012) tractography was run on the normalized white matter FOD with the probabilistic algorithm iFOD2(J.-D. Tournier et al., 2010) and a dynamic seeding strategy(R. E. Smith et al., 2015) to generate tractograms of 3 million streamlines. All tractograms were filtered in a multi-stage process (see(Lu et al., 2025) for more detail):

1. Any erroneous streamlines that failed to connect two gray matter ROIs were manually removed to ensure they did not bias the model in step 2.
2. COMMIT v2.1(Daducci et al., 2013, 2015) (convex optimization modeling for microstructure informed tractography) was run on the stage-1 filtered tractograms. COMMIT evaluates the contribution of each streamline to the measured diffusion signal and assigns weights that best reconstruct the voxel-wise diffusion profile. Streamlines which are not supported by the underlying microstructure are down-weighted or eliminated. COMMIT was optimized to the DWI image using a Stick-Zeppelin-Ball forward model with diffusivities Ð_∥_ = 1.7e^-3^, Ð_⊥_ = 0.51e^-3^ and Ð_𝑖𝑠𝑜_ = 1.7e^-3^, 3.0e^-3^ mm^2^/s (gray matter and CSF). Estimates of the signal fraction or intra-cellular volume fraction (ICVF) of each streamline were extracted from the COMMIT model. Streamlines < 1e^-12^ were filtered as they did not contribute to the global DWI signal. A volumetric ICVF map was also obtained from COMMIT. This was used to calibrate the MT_sat_ map yielding a myelin volume fraction (MVF) map as:

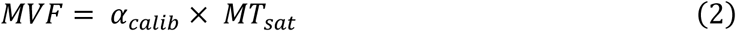

where 𝛼_calib_ is the global calibration factor as described in(Lu et al., 2025).

3. The stage-2 filtered tractograms were optimized to the MVF map with a myelin-based COMMIT extension(Schiavi et al., 2022) yielding a signal fraction for each streamline which corresponded to a myelin cross-sectional area. Streamlines < 1e^-12^ were filtered as they did not contribute to the global MVF signal.
4. To get the final signal fraction (ICVF) for the surviving streamlines, a modified version of step 2 was repeated on the stage-3 filtered tractograms. Instead of optimizing COMMIT to the DWI signal itself, a scaled version was computed (1-MVF), which was then normalized by the b = 0 images. This was done to account for the potential bias of myelination to the ICVF estimates from COMMIT. A very small number of streamlines < 1e^-12^ were filtered in this stage as well.

These filtered tractograms were used for all structural connectomes i.e., both the underlying streamlines and edge connection density were uniform across all variants of weighted structural connectomes. Although our filtering pipeline made it possible to estimate a connectome weighted by tract-specific MVF, such a myelin metric is highly collinear with edge caliber (derived from COMMIT in our analysis), which would represent a significant confound in our multiple regression framework. Thus, these were not used in this study. Instead, we are estimating edge myelin density using MTsat tractometry, which is more complementary to edge caliber (Figure S10).

#### Weighted structural connectomes

The edge *caliber* connectome was computed by summing the final COMMIT weights (signal fractions) of all remaining streamlines connecting each node pair normalized by total node volume for each pair. The edge *myelin* connectome was computed by applying tractometry(S Bells et al., 2011) to the MTsat image: the median MTsat value was computed along each streamline, followed by the average across streamlines for each node pair. The edge *length* connectome was computed as the mean length of streamlines for each node pair. The Schaefer-400(Schaefer et al., 2018) cortical atlas was used to define network nodes in all primary analyses.

#### Group-consensus structural connectomes

We developed a group averaging method through rigorous testing to maximally preserve subject-level features. This was carried out in a two-stage process. (1) Consistency-based filtering(Roberts et al., 2017) targeted to cross-subject variance in number of streamlines edge weights was used to filter all subject-level structural connectomes to a threshold of 30% (the final density if averaging was performed at this stage). (2) Distance-dependent thresholding(Betzel et al., 2019) was then applied to generate group-representative structural connectomes which preserved subject-level within- and between-hemisphere edge length distributions. Final group-level connection density was 23%, which was approximately the median subject-level density.

### Functional connectivity network reconstruction

Functional connectivity was computed as the zero-lag Pearson cross-correlation of fMRI time series. These values were Fisher z-transformed and averaged across subjects yielding a single in-sample-fMRI functional connectome in the Schaefer-400 parcellation.

#### HCP data

Fully-processed fMRI and MEG functional connectivity matrices were obtained at the group-level in the Schaefer-400 parcellation. These connectomes were derived from resting-state data acquired in healthy young adults (n = 33; age range 22 to 35 years) as part of the Human Connectome Project (HCP; S900 release(Van Essen et al., 2013)). In a previous study(Shafiei et al., 2022), these specific datasets were selected from the small proportion of the full cohort which contained MEG data for unrelated subjects, and we elected to reuse these same data as the number of participants and their age range matched closely to our in-house data sample. The MEG data corresponded to 6 minutes of scanning (sampling rate = 2,034.5 Hz; anti-aliasing low-pass filter = 400 Hz), and the fMRI data were comprised of 4 scans of 15 minutes across 2 sessions (TR = 720 ms; 3 Tesla; 2 mm isotropic). Static functional connectivity was computed as (1) the Pearson cross-correlation of fMRI time series and (2) the amplitude envelope correlation(Bruns et al., 2000) of MEG time series data. An orthogonalization process was then applied to correct for spatial leakage in the MEG functional connectomes(G. L. Colclough et al., 2015). Group-averaging yielded one fMRI and six MEG functional connectomes corresponding to the canonical electrophysiological frequency bands: delta (δ: 2-4 Hz), theta (θ: 5-7 Hz), alpha (α: 8-12 Hz), beta (β: 15-29 Hz), low gamma (γ_lo_: 30-59 Hz), and high gamma (γ_hi_: 60-90Hz). See(Shafiei et al., 2022) for full processing details.

### Multi-linear modeling

We quantified the role of white matter features in predicting synchronized brain function using multiple linear regression. The dependent variable (response variable) was functional connectivity (FC) from one of the following modalities: in-sample-fMRI (BOLD_in_), HCP-fMRI (BOLD_out_) or HCP-MEG in the 6 canonical frequency bands (n = 8 unique dependent variables). The predictors were structural connectivity (SC) weights quantifying the *caliber*, *myelin* and *length* of each edge. Interaction terms were included for both caliber and length with myelin.

Only edges with SC ≠ 0 were included. The *adjusted R^2^*(coefficient of determination) was interpreted as the magnitude of structure-function coupling. The contribution of each predictor was quantified with dominance analysis(Azen & Budescu, 2003; Budescu, 1993), which measures relative predictor importance as the incremental change in R^2^ from the addition (or absence) of a given predictor to all possible subset models. Dominance analysis was adapted to handle interaction terms for this work. Variables with an absolute skewness > 1.5 were log-transformed, and all variables were z-scored. Modeling was restricted to group-level data.

Parametric t-tests were used to exclude any ß-coefficients with p > 0.05.

**Global model**. The whole-brain linear regression model was computed as:

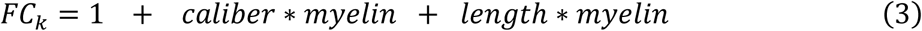

where the predictors *caliber*, *myelin* and *length* are the vectorized lower triangle elements of the corresponding weighted structural connectome, and the dependent variable *FC_k_* is the corresponding vectorized lower triangle elements of the *k*^th^ FC dataset.

#### Network model

Nodes (cortical parcels) were grouped by affiliation to the 7-canonical resting state sub-networks(Thomas Yeo et al., 2011), and linear regression was performed within pairwise combinations of sub-networks as:

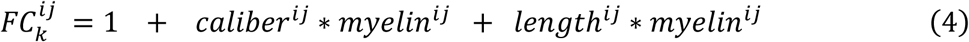

where the FC and SC data corresponded to the weights of network edges linking the nodes of subnetworks *i* and *j*. Within-subnetwork modeling is quantified by the case where *i* = *j*.

**Node model**. Linear regression was performed separately for each node as:

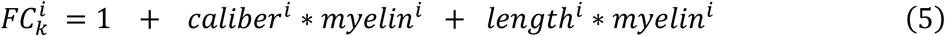

where the FC and SC data corresponded to the weights of network edges linking node *i* to all other *j* ≠ *i* nodes.

### Null models

To account for the lack of strictness of parametric t-tests applied to our data, we additionally assessed the significance of our results using edgewise permutation testing with 1000 repetitions. When testing the model outputs, the FC edge weights were shuffled to ensure the relationships between the structural predictors were preserved. For the global model, a spatial constraint was added to the permutation test by shuffling FC edge weights within 10 equally sized bins defined by Euclidean distance. One-sided p-values were computed in which the empirical value was included in the null distribution to avoid p_perm_ = 0.

## Data availability

All data necessary to replicate this work are freely available at https://github.com/TardifLab/Modeling-Myelin-FC.

## Code availability

All code necessary to replicate this work are freely available at https://github.com/TardifLab/Modeling-Myelin-FC.

## Acknowledgments

The authors acknowledge research support from the National Science and Engineering Research Council of Canada (NSERC), the Canadian Institutes for Health Research (CIHR), Fonds de recherche du Québec – Santé (FRQ-S), CFREF Healthy Brains for Healthy Lives, the Killam Trusts, and Brain Canada.

