## supplemental figures for "The role of white matter myelin in structural-functional network coupling"

### Supplementary Information

Mark C. Nelson[1,2], Wen Da Lu[2,3], Ilana R. Leppert[2], Heather A. Hansen[1], Christopher D. Rowley[4], Bratislav Misic[1,2], and Christine L. Tardif [1,2,3]

[1]*Department of Neurology and Neurosurgery, McGill university, Montreal, QC, Canada.* [2]*McConnell Brain Imaging Centre, Montreal Neurological Institute and Hospital, Montreal, QC, Canada.* [3]*Department of Biomedical Engineering, McGill University, Montreal, QC, Canada.* [4]*Department of Physics and Astronomy, McMaster University, Hamilton, ON, Canada.*

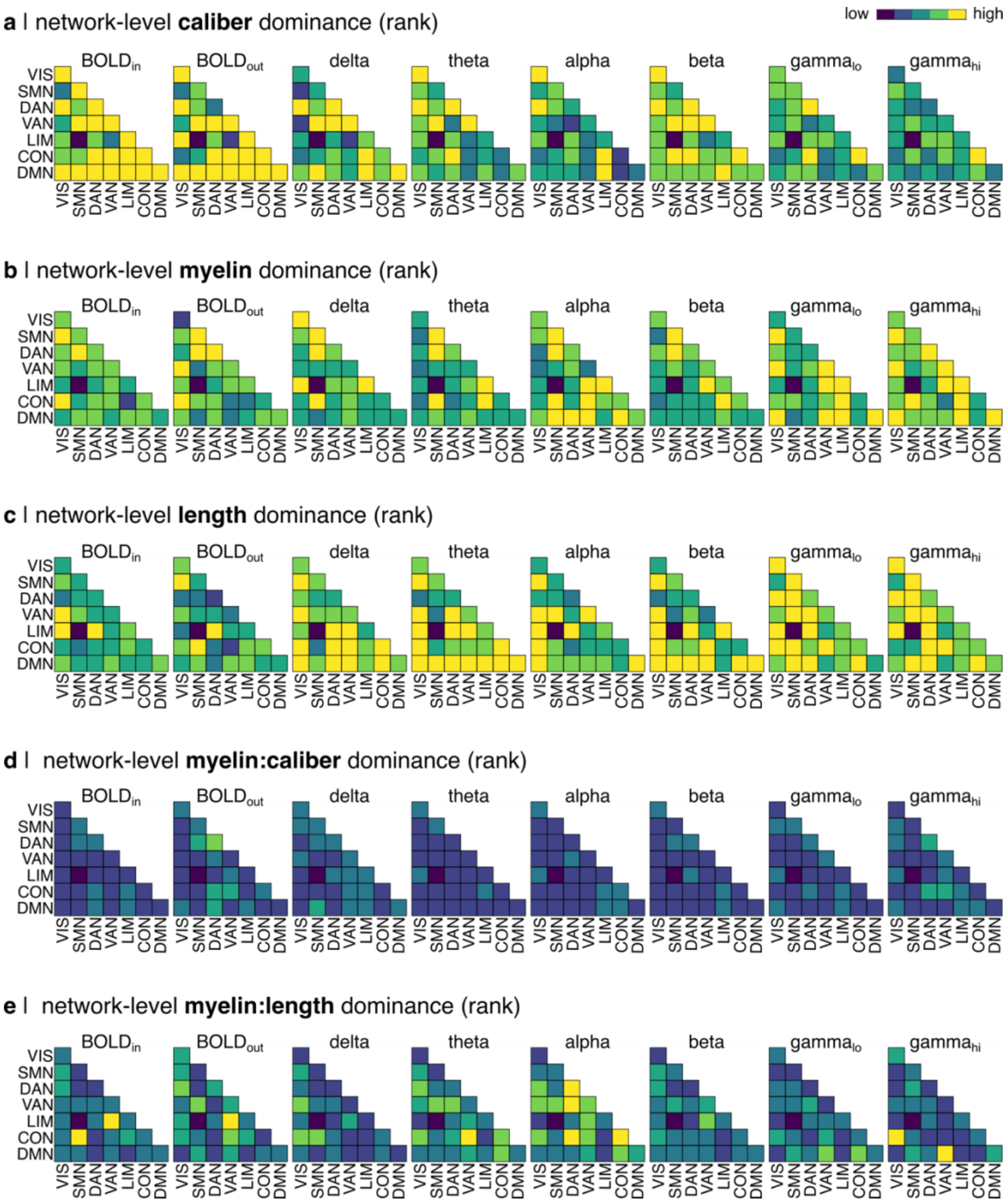


*Figure S1. Dominance rankings for **(a)** caliber, **(b)** myelin, **(c)** length, (d) the interaction of the myelin and caliber, and (e) the interaction of myelin and length for every FC model at the resting-state network-level. The canonical Yeo-7-Networks are used: visual (VIS), somatomotor (SMN), dorsal attention (DAN), salience ventral attention (VAN), limbic (LIM), fronto-parietal control (CON), default mode (DMN).*

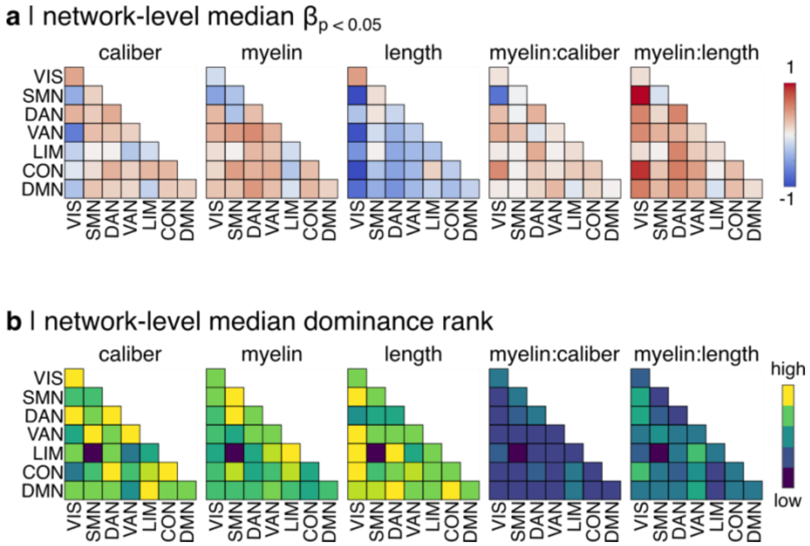


*Figure S2. Summary of the role of structural features in predicting FC at the resting-state network level. Median values are computed across FC networks for* ***(a)*** *standardized ß-coefficients and* ***(b)*** *dominance*

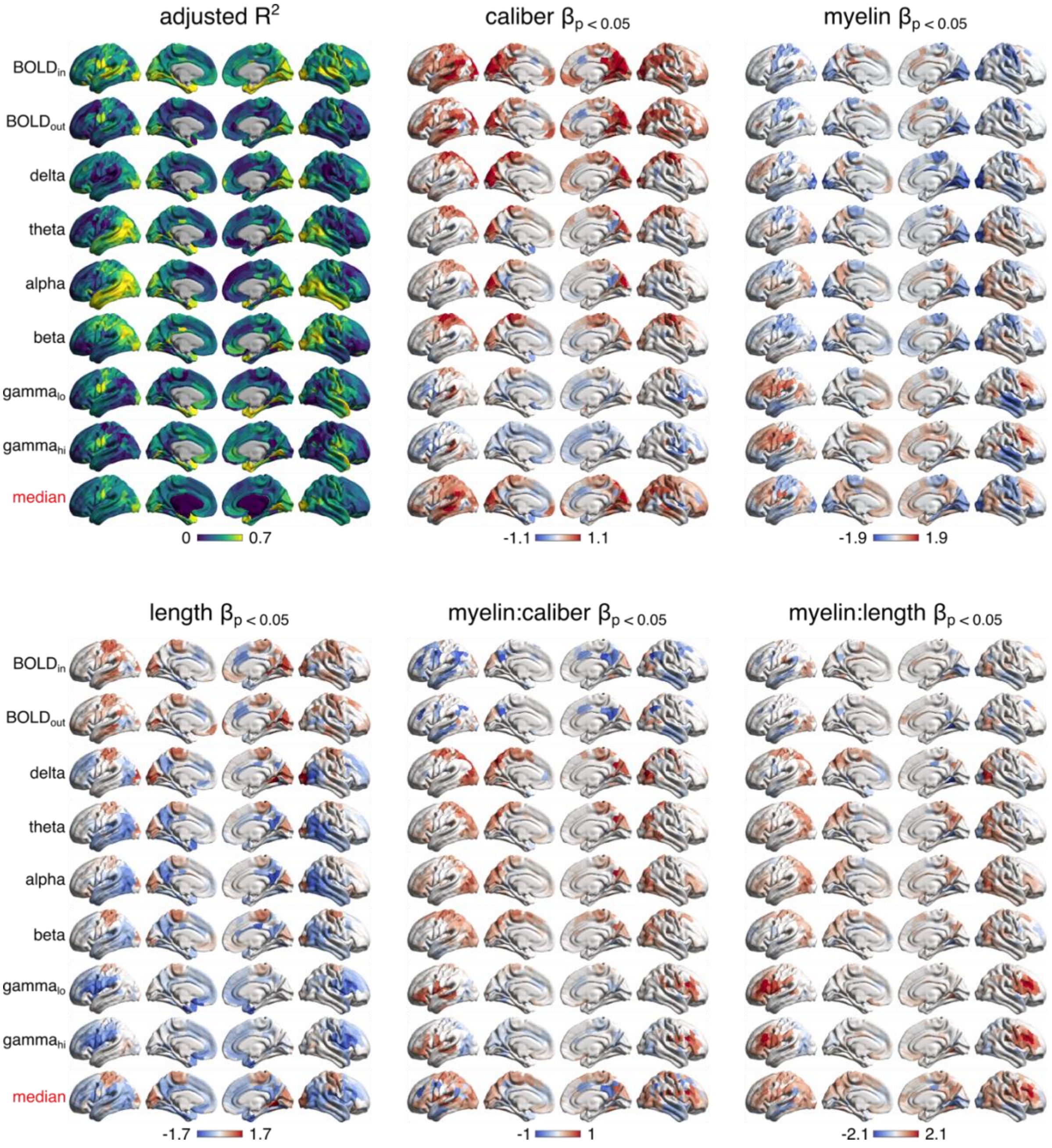


*Figure S3. Node-level SFC (adjusted $R^2$) and β-coefficients for each FC network. The median across FC networks is shown for each node on the bottom row. Only significant β-coefficients are included (t-test, $p < 0.05$).*

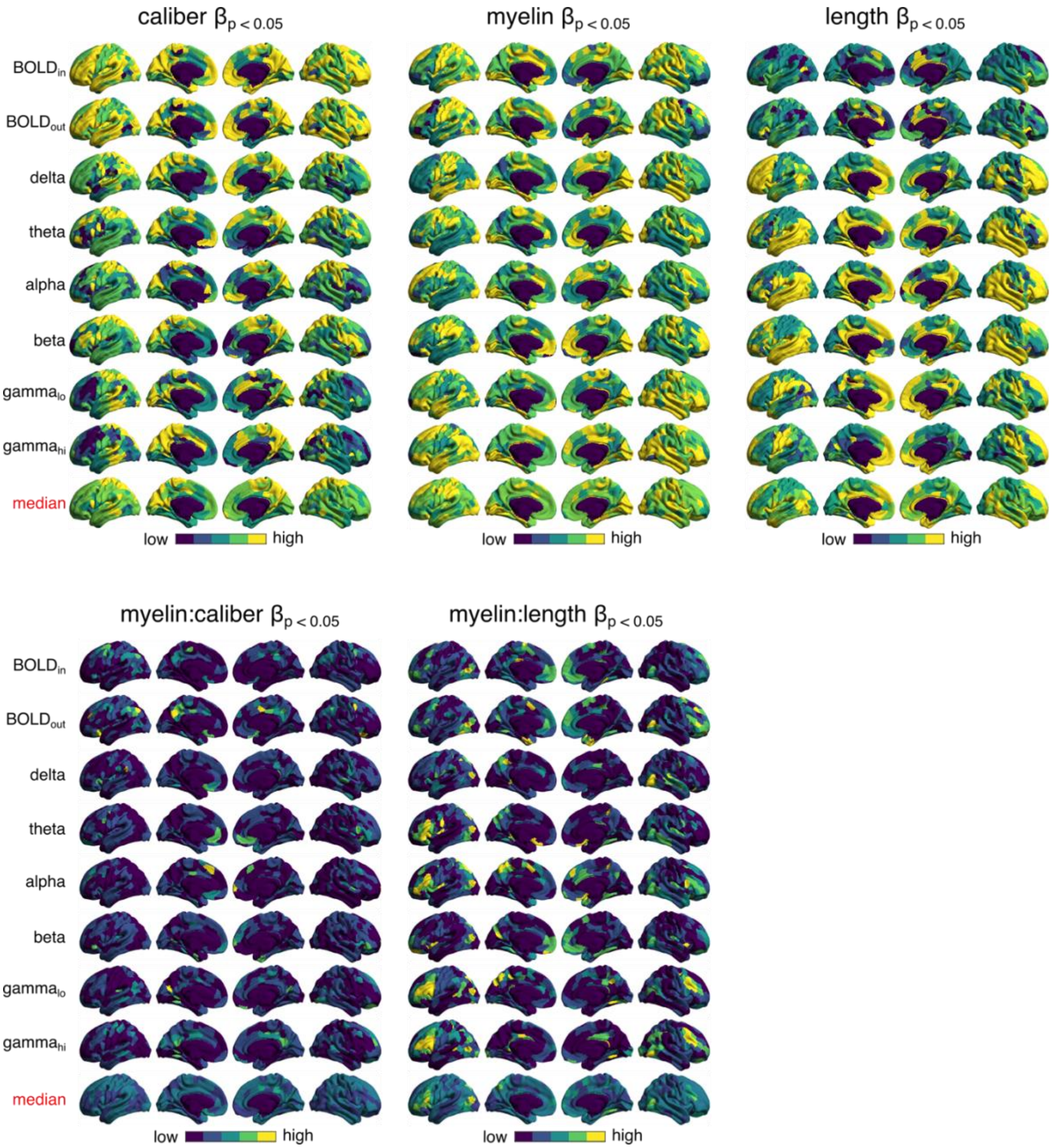


*Figure S4. Dominance rankings for each structural feature are shown for every FC model at the node-level. The median across FC networks is shown for each node on the bottom row.*

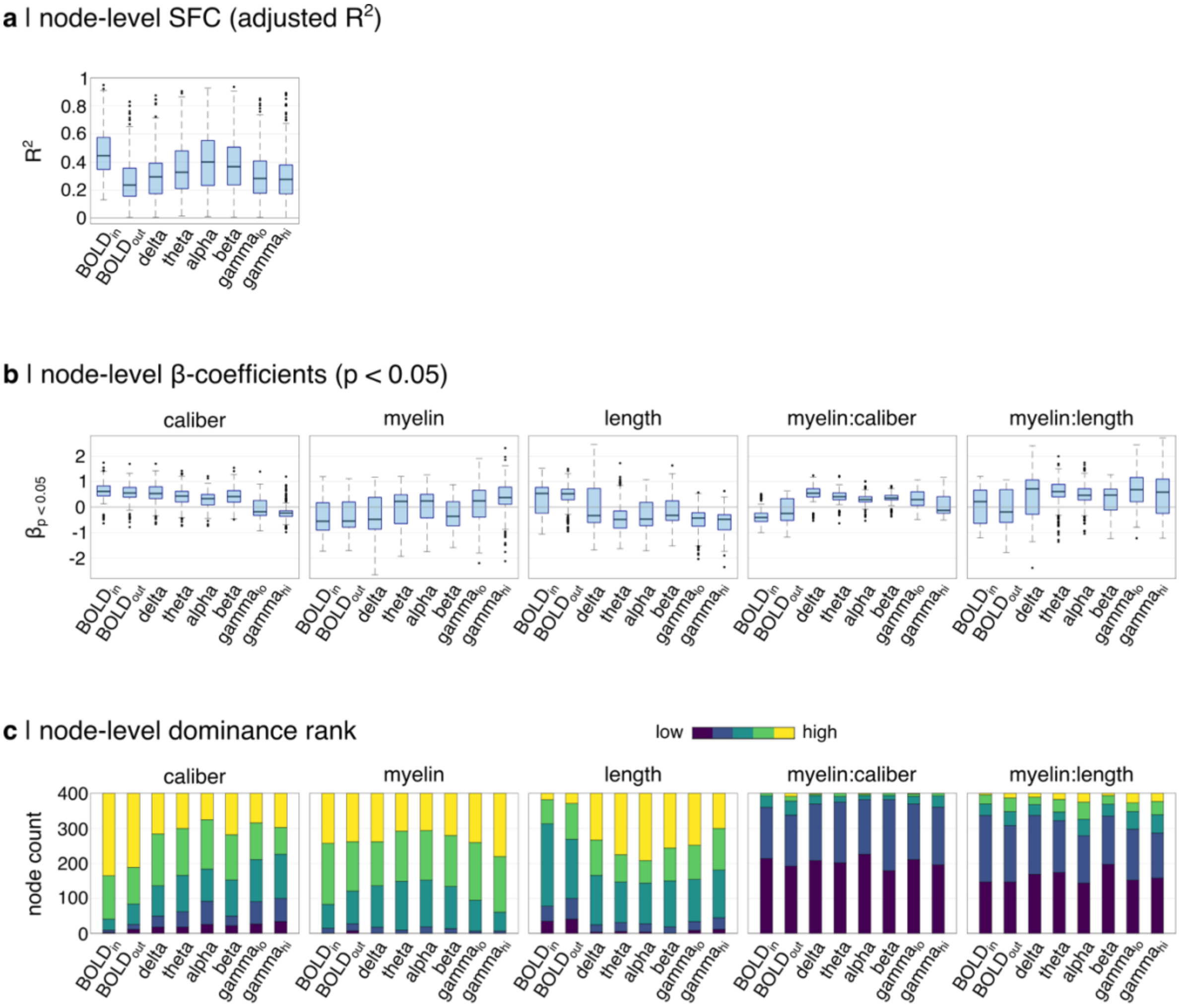


*Figure S5. Node-level results of the full model with interactions is summarized for every FC dataset. The distributions of **(a)** SFC (adjusted $R^2$) and **(b)** standardized β-coefficients are shown, as well as **(c)** the count of dominance rankings.*

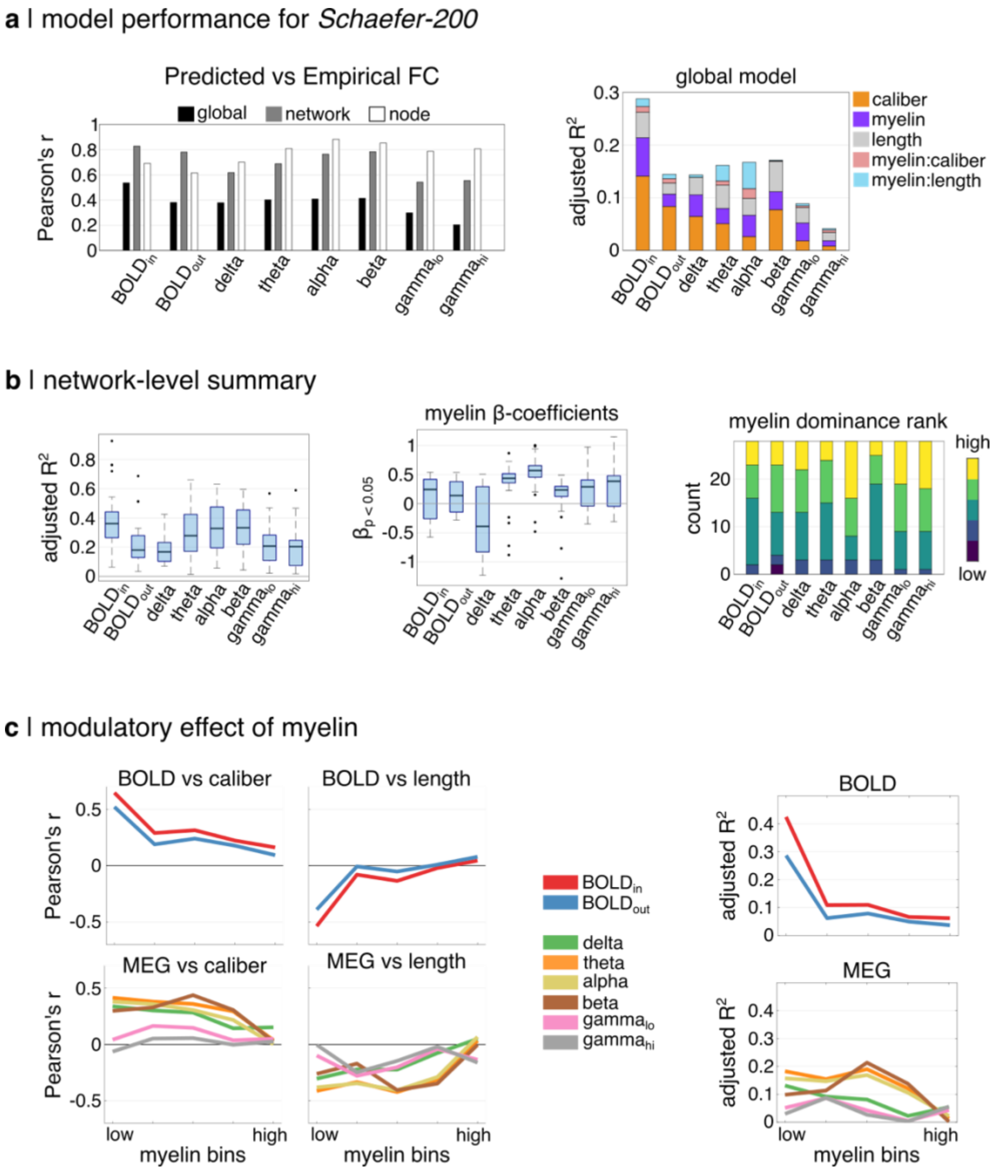


*Figure S6. Replication using Schaefer-200 parcellation. Processing steps were identical, except consistency-based filtering threshold was set to 35% (vs 30% for Schaefer-400). Overall modeling trends are highly similar to the main analysis.*

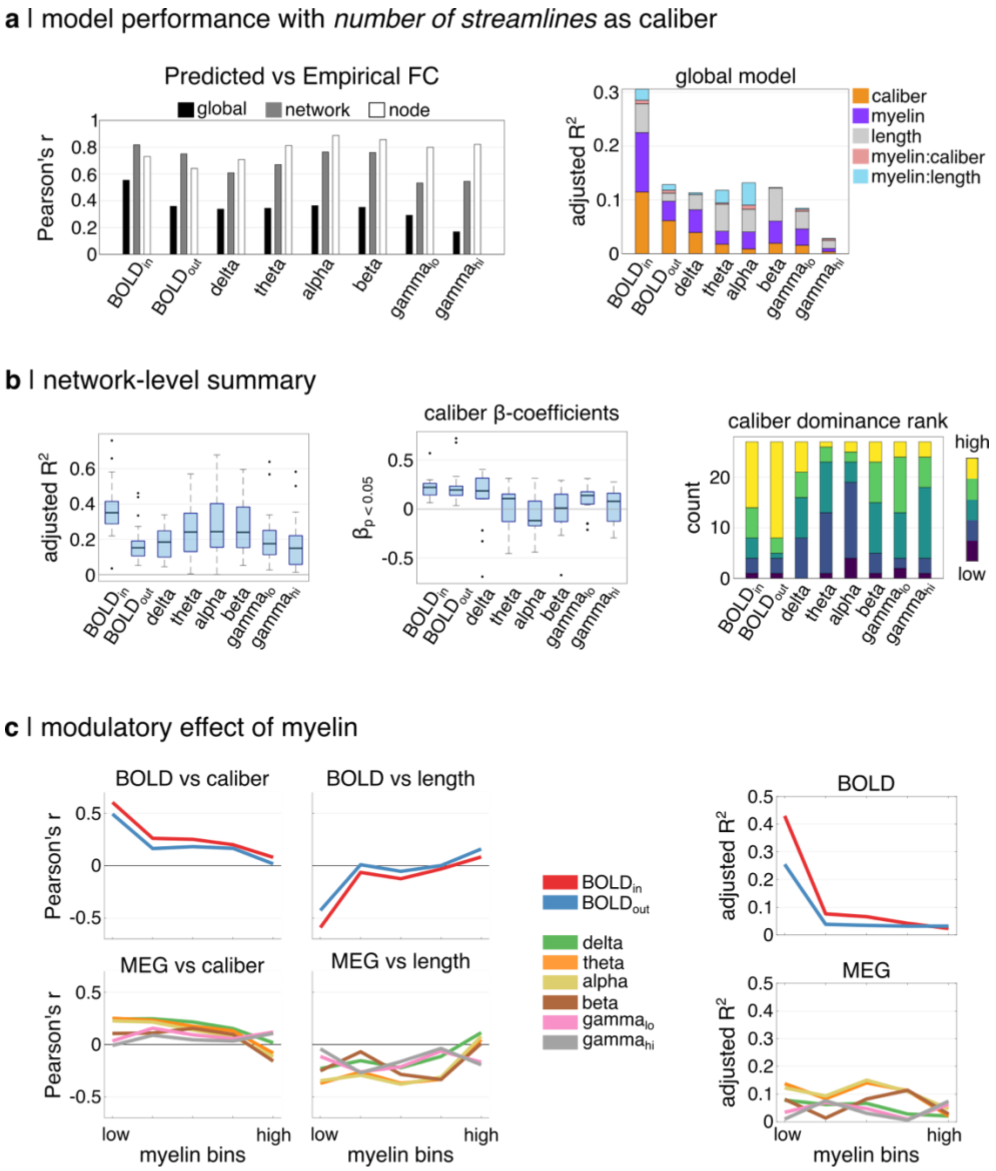


*Figure S7. Replication using streamline counts as an estimate of edge caliber. All other data and processing steps are identical to the main analysis. Overall modeling trends are highly similar to the main analysis. Streamline counts are generally a weaker predictor of FC relative to COMMIT.*

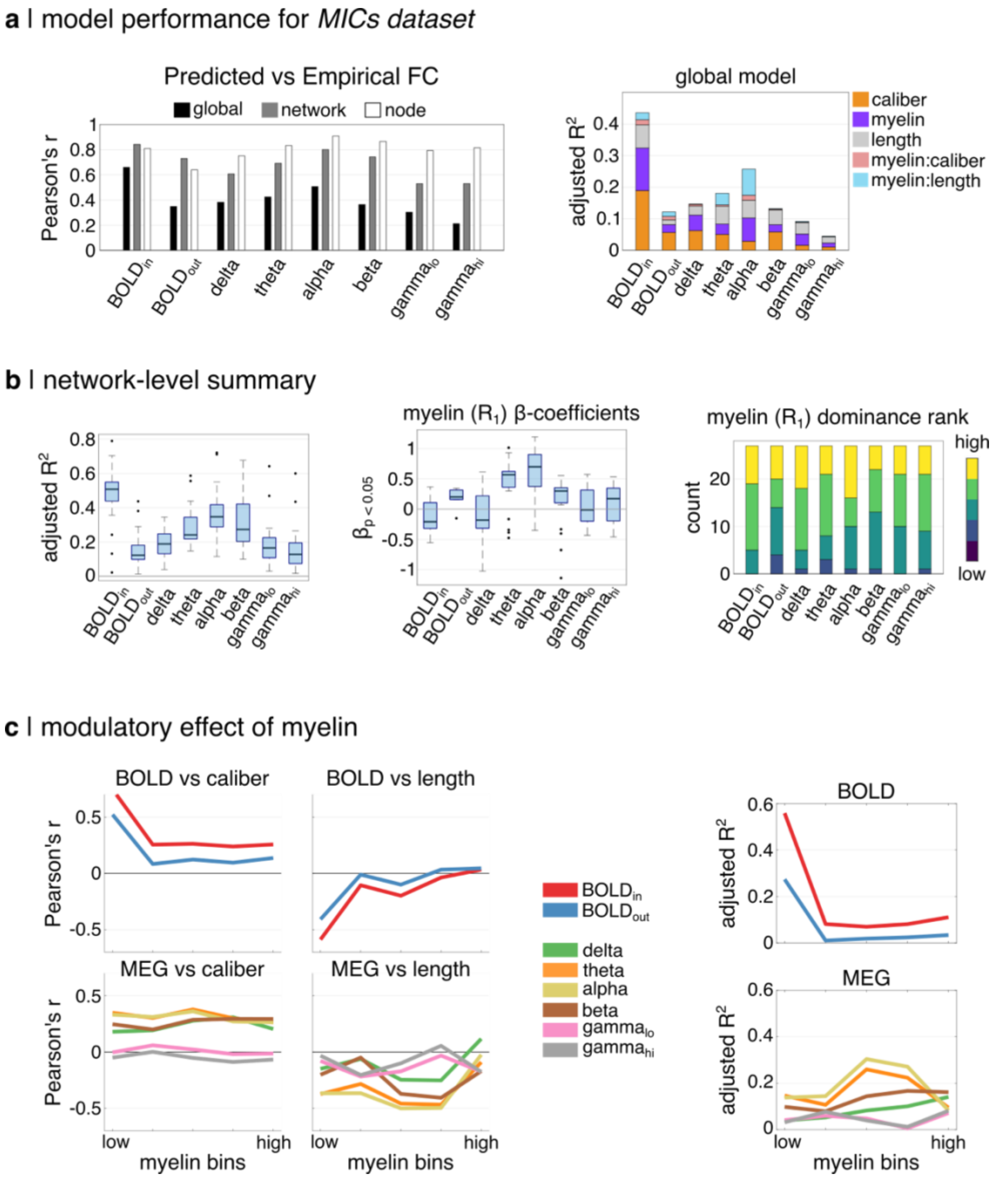


*Figure S8. Replication using an independent multi-modal MRI dataset. Edge myelin was quantified using the longitudinal relaxation rate ($R_1$). Edge caliber was quantified using COMMIT. Overall modeling trends are highly similar to the main analysis. $R_1$-based myelin ranks higher in predictor dominance across network pairs relative to $MT_{sat}$. These data and their associated publication are available here [https://portal.conp.ca/dataset?id=projects/mica-mics](https://portal.conp.ca/dataset?id=projects/mica-mics) and [https://doi.org/10.1038/s41597-022-01682-y](https://doi.org/10.1038/s41597-022-01682-y), respectively.*

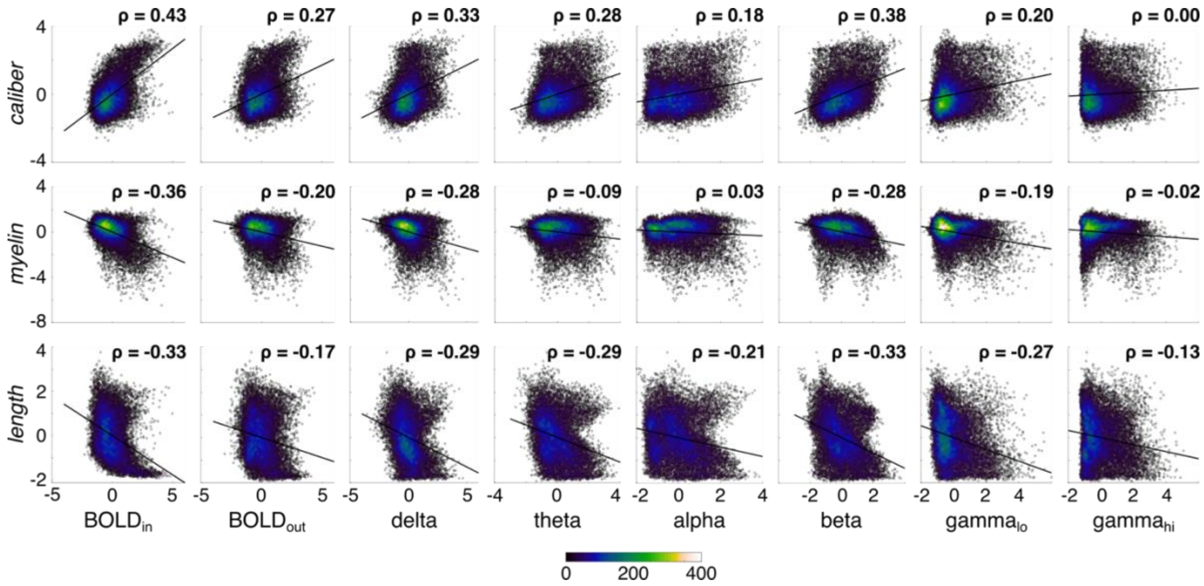


*Figure S9. Interrelationships of structural features and FC networks. Spearman's rank correlations, and a best fit line are shown for each plot. Color represents data density. Edge caliber is log-transformed, and all data is z-scored.*

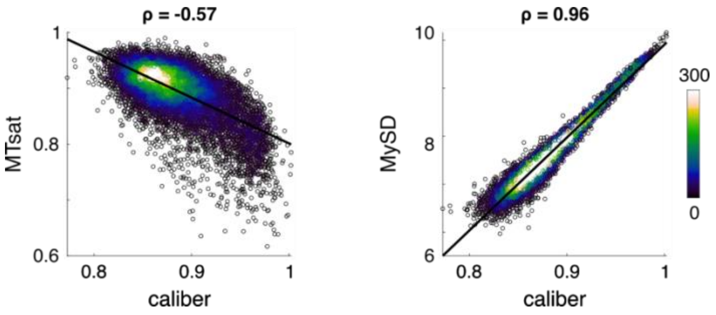


*Figure S10. Spearman's rank correlation of myelin metrics with edge caliber.*
